# SARS-CoV-2 receptor Angiotensin I-Converting Enzyme type 2 (ACE2) is expressed in human pancreatic β-cells and in the human pancreas microvasculature

**DOI:** 10.1101/2020.07.23.208041

**Authors:** Daniela Fignani, Giada Licata, Noemi Brusco, Laura Nigi, Giuseppina E. Grieco, Lorella Marselli, Lut Overbergh, Conny Gysemans, Maikel L. Colli, Piero Marchetti, Chantal Mathieu, Decio L. Eizirik, Guido Sebastiani, Francesco Dotta

**Affiliations:** Diabetes Unit, Department of Medicine, Surgery and Neurosciences, University of Siena, Siena, Italy; Fondazione Umberto Di Mario, c/o Toscana Life Sciences, Siena, Italy; Tuscany Centre for Precision Medicine (CReMeP), Siena, Italy; Department of Clinical and Experimental Medicine, University of Pisa, Pisa, Italy; Clinical and Experimental Endocrinology (CEE), Katholieke Universiteit Leuven (KULEUVEN), Leuven, Belgium; ULB Center for Diabetes Research, Medical Faculty, Université Libre de Bruxelles, Brussels, Belgium; Indiana Biosciences Research Institute, Indianapolis, Indiana, USA

**Keywords:** Diabetes, COVID-19, ACE2, beta-cell, human pancreatic islets, SARS-CoV-2, inflammation

## Abstract

Increasing evidence demonstrated that the expression of Angiotensin I-Converting Enzyme type 2 (ACE2), is a necessary step for SARS-CoV-2 infection permissiveness. In the light of the recent data highlighting an association between COVID-19 and diabetes, a detailed analysis aimed at evaluating ACE2 expression pattern distribution in human pancreas is still lacking. Here, we took advantage of INNODIA network EUnPOD biobank collection to thoroughly analyse ACE2, both at mRNA and protein level, in multiple human pancreatic tissues and using several methodologies.

Using multiple reagents and antibodies, we showed that ACE2 is expressed in human pancreatic islets, where it is preferentially expressed in subsets of insulin producing β-cells. ACE2 is also is highly expressed in pancreas microvasculature pericytes and moderately expressed in rare scattered ductal cells. By using different ACE2 antibodies we showed that a recently described short-ACE2 isoform is also prevalently expressed in human β-cells.

Finally, using RT-qPCR, RNA-seq and High-Content imaging screening analysis, we demonstrated that pro-inflammatory cytokines, but not palmitate, increases ACE2 expression in the β-cell line EndoC-βH1 and in primary human pancreatic islets.

Taken together, our data indicate a potential link between SARS-CoV-2 and diabetes through putative infection of pancreatic microvasculature and/or ductal cells and/or through direct β-cell virus tropism.

## 1 Introduction

The expression of molecules that act as receptors for viruses determine tissue-specific tropism. SARS-coronavirus 2 (SARS-CoV-2), that leads to the respiratory illness coronavirus disease 2019 (COVID-19), uses its surface envelope Spike glycoprotein (S-protein) to interact and gain access to host cells through the Angiotensin-I converting enzyme-2 (ACE2) receptor. As such, S-protein-ACE2 binding is the key determinant for virus entry, propagation and transmissibility of COVID-19-related disease (1,2).

Artificially induced ACE2 *de-novo* expression in ACE2-negative cell lines is a necessary step to SARS-CoV and SARS-CoV-2 infection (3,4). SARS-CoV-2 does not enter cells that do not express ACE2 and does not use other coronaviruses receptors, such as aminopeptidase N (APN) and dipeptidyl peptidase 4 (DPP4), thus being fully dependent on ACE2 presence in host cells (5). Additional host co-factors, such as transmembrane protease TMPRSS2, cathepsin B/L and furin protease, have been shown to enhance efficiency of SARS-CoV-2 cell entry by processing the S-protein and eliciting membrane fusion and syncytia formation (6). The central role played by ACE2 in SARS-CoV-2 infection has been further supported by evidence that SARS-CoV-2 infection is driven by ACE2 expression level (6). SARS-CoV-2 mainly targets cells of the nasal, bronchial and lung epithelium, causing respiratory-related symptoms; however, growing evidence shows that other tissues can also be infected.

Several reports indicate a wide, although variable, distribution of ACE2 expression patterns among different tissues (7–9), thus underlining a potential different virus infection susceptibility among cell types. The fact that COVID-19 disease may lead to multiple organ failure (10,11) shows the crucial relevance for understanding the molecular mechanisms of host cell factors used by SARS-CoV-2 to infect their target tissues.

Recent studies showed that older adults and those with chronic medical conditions like heart and lung disease and/or diabetes mellitus are at the highest risk for complications from SARS-CoV-2 infection. Of importance, a yet unresolved conundrum relies on the recently hypothesized bidirectional relationship between COVID-19 and diabetes mellitus (12,13). This concept is supported by reports in which impaired glycaemic control is associated with increased risk of severe COVID-19. Indeed, elevated blood glucose concentrations and deterioration of glycaemic control may contribute to increased inflammatory response, to abnormalities in the coagulation system and to impairment of ventilatory function, thus leading to severe COVID-19 disease and to a worse prognosis (14). Interestingly, acute hyperglycaemia has been observed at admission in a substantial percentage of SARS-CoV-2 infected subjects, regardless of the past medical history of diabetes (15–18). The same observations were previously made in SARS-CoV-1 pneumonia during 2003 SARS epidemic (19).

A recently published case report described autoantibody-negative insulin-dependent diabetes onset in a young patient who was infected by SARS-CoV-2 seven weeks before diabetes symptoms occurrence (20). Additional previous studies further support such observation(21,22). This indicates the possibility of a link between SARS-CoV-2 infection and new-onset diabetes through potential direct infection of pancreatic islets or additional indirect mechanisms. Indeed, an *in-vitro* infection model of human pluripotent stem cells derived β-cells exposed to SARS-CoV-2 (23) showed permissiveness of these pre-β-cells to the virus. However, whether fully mature primary beta-cells or other cells of the human pancreas are indeed permissive to SARS-CoV-2 infection remains to be clarified.

To address this question, we screened the ACE2 expression pattern in human pancreata obtained from adult non-diabetic multiorgan donors and in the insulin-producing human β-cell line EndoC-βH1, using different methodologies, multiple reagents, and publicly available or in-house generated RNA sequencing datasets. Our data indicate that ACE2 is expressed by pancreas microvasculature, by scattered ductal cells and by a subset of human β-cells. These different cell types are thus potentially prone to SARS-CoV-2 infection. We also identified a differential distribution of the two recently discovered ACE2 isoforms (24,25). Exposure of EndoC-βH1 human beta-cell line and human pancreatic islets to pro-inflammatory cytokines significantly increased ACE2 expression. Taken together, our data suggest a potential link between SARS-CoV-2 infection and new onset diabetes, which deserves further investigation based on long-term follow up of patients recovered from COVID-19 disease.

## 2 Materials and Methods

### 2.1 Human donors

Human pancreatic sections analysed in this study were obtained from pancreata of brain-dead adult non-diabetic multiorgan donors within the European Network for Pancreatic Organ Donors with Diabetes (EUnPOD), a project launched in the context of the INNODIA consortium (www.innodia.eu). Whole pancreata were processed following standardized procedures at University of Pisa. Formalin fixed paraffin embedded (FFPE) pancreatic tissue sections and frozen OCT pancreatic tissue sections were obtained from n=7 adult non-diabetic multiorgan donors, and from n=1 longstanding T1D donor pancreas (**Table S1**). In INNODIA EUnPOD network, pancreata not suitable for organ transplantation were obtained with informed written consent by organ donors’ next-of -kin and processed with the approval of the local ethics committee of the Pisa University.

### 2.2 Human pancreatic islets

Human pancreatic islets were obtained from n=4 non-diabetic multi-organ donors (**Table S1**). Briefly, purified islets were prepared by intraductal collagenase solution injection and density gradient purification, as previously described (26). At the end of the isolation procedure, fresh human pancreatic islets preparations were resuspended in CMRL culture medium (cat. 11-530-037, ThermoFisher Scientific, Waltham, MA, USA) supplemented with L-Glutamine 1% (cat.G7513-100ML), Antibiotic/Antimicotic 1% (A5955-100ML, Sigma Aldrich, St. Louis, MO, USA), FBS 10% and cultured at 28°C in a 5% CO_2_ incubator.

### 2.3 Cell culture

EndoC-βH1 human β-cell line (27,28) was obtained by UniverCell-Biosolutions (Toulouse-France) and used for all experiments between passages 78-88. EndoC-βH1 were cultured at 37 °C with 5% CO_2_ in coated flask (coating medium composition: DMEM high-glucose cat. 51441C, Penicillin/Streptomycin 1% cat. P0781, ECM 1% cat. E1270 and Fibronectin from bovine plasma 0.2% cat. F1141 - all from Sigma Aldrich, St. Louis, MO, USA) and maintained in culture in low-glucose DMEM (cat. D6046) supplemented with 2% BSA fraction V (cat. 10775835001), β-Mercaptoethanol 50 μM (cat. M7522), L-Glutamine 1% (cat. G7513), Penicillin/Streptomycin 1% (cat. P0781), Nicotinamide 10 mM (cat. N0636), Transferrin 5.5 μg/mL (cat. T8158) and Sodium selenite 6.7 ng/mL (cat. S5261) (all from Sigma Aldrich, St. Louis, MO, USA).

HeLa cells (ATCC CCL-2), passages 33-34, were cultured at 37 °C with 5% CO_2_ in a 100 mm petri plate and maintained in culture in high glucose DMEM (cat. 51441C) supplemented with L-Glutamine 1% (cat. G7513), Antibiotic/Antimycotic 1% (A5955-100ML, Sigma Aldrich, St. Louis, MO, USA) and FBS 10%.

In order to evaluate ACE2 expression in human β cells under diabetogenic stress conditions, EndoC-βH1 cell line was subjected to palmitate-induced lipotoxic and inflammatory stress. Briefly, EndoC-βH1 cells were plated at a density of 2,5×10^5^/well in 24-well plates or 5×10^4^/well in 96-well plates. After 48 h, palmitate and inflammatory stimuli were performed as previously described (29,30). In details, palmitate and inflammatory stresses have been induced respectively by 2 mM of Sodium Palmitate (cat. P9767-5G - Sigma Aldrich, St. Louis, MO, USA) or 0,5% EtOH (as control treatment) for 24 h, or cytokines mix IL-1β (50 U/mL) (cat. #201-LB-005 - R&D System, Minneapolis, MN, USA), TNFα (1000 U/mL) (cat. T7539 - Sigma Aldrich, St. Louis, MO, USA) and IFNγ (1000 U/mL) (cat. 11040596001-Roche, Basilea, Switzerland) for 24 h.

### 2.4 Laser capture microdissection (LCM)

Pancreatic human tissue samples (n=5) from EUnPOD multiorgan donors (**Table S1**) were frozen in Tissue-Tek OCT compound and then 7-μm thick sections were cut from frozen O.C.T. blocks. Sections were fixed in 70% ethanol for 30 s, dehydrated in 100% ethanol for 1 min, in 100% ethanol for 1 min, in xylene for 5 min and finally air-dried for 5 min. Laser capture microdissection (LCM) was performed using an Arcturus XT Laser-Capture Microdissection system (Arcturus Engineering, Mountain View, CA, USA) by melting thermoplastic films mounted on transparent LCM caps (cat.LCM0214 - ThermoFisher Scientific, Waltham, MA, USA) on specific islet areas. Human pancreatic islets were subsequently visualized through islet autofluorescence for LCM procedure. Thermoplastic films containing microdissected cells were incubated with 10 μl of extraction buffer (cat. kit0204 - ThermoFisher Scientific, Waltham, MA, USA) for 30 min at 42 °C and kept at −80°C until RNA extraction. Each microdissection was performed within 30 min from the staining procedure. Overall n=50 microdissected pancreatic islets from each case were analysed.

### 2.5 RNA extraction from LCM isolated human pancreatic islets

Total RNA was extracted from each LCM sample using PicoPure RNA isolation kit Arcturus (cat. kit0204 - ThermoFisher Scientific, Waltham, MA, USA) following manufacturer’s procedure. Briefly, the cellular extracts were mixed with 12.5 μl of EtOH (100%) and transferred to the purification column filter membrane. DNase treatment was performed using RNase-Free DNase Set (cat. 79254 - Qiagen, Hilden, Germany). Total RNA was eluted in 11 μl of DNase/RNase-Free Water and LCM captures deriving from human sample were pooled and subjected to a subsequent concentration through Savant SpeedVac SC100 centrifugal evaporator. Agilent 2100 Bioanalyzer technology with RNA Pico chips (cat. 5067-1513 Agilent Technologies, Santa Clara, CA, USA) was performed for each RNA sample, in order to analyse RNA integrity (RIN) and concentration.

### 2.6 RNA extraction from cells and tissues

For gene expression evaluation, total RNA was extracted from approximately 3.0×10^5^ EndoC-βH1 or from fresh lung tissue (0.5×0.5×0.5cm), obtained from a lung tumor surgery donor by dissecting a not affected portion of the tissue (obtained with informed written consent of the patient and approved by the local Ethics Committee at the University of Siena). Direct-zol RNA Miniprep Kit (cat. R202-Zymo Research, Irvine, CA, US) was adopted following manufacturer’s instructions. Briefly, the pelleted cells were resuspended in QIAzol (cat. 79306, Qiagen), mixed with equal volume of Ethanol 100% and transferred to Zymo-Spin™ IICR Column. DNase digestion was performed using RNase-Free DNase Set (cat. 79254). RNA was eluted in 30 μl of DNase/RNase-Free Water. Fresh lung tissue was maintained in PBS1X on ice immediately after surgery, until ready for the RNA extraction. Homogenization of the tissue was performed using 600 μl of QIAzol and Lysing Matrix latex beads (MP Biomedicals, cat. 6913-100) in FastPrep-24 automated homogenizer (1min, full speed). The homogenate was diluted with vol/vol Ethanol 100% and then transferred in Zymo-Spin™ IICR Column. RNA extraction was performed following manufacturer’s instructions.

### 2.7 RT-Real Time PCR analysis

Total RNA extracted from EndoC-βH1 and collagenase-isolated human pancreatic islets samples was quantified using Qubit 3000 Fluorometer (ThermoFisher Scientific, Waltham, MA, USA), while those extracted from LCM-islets were quantified using 2100 Bioanalyzer-RNA 6000 Pico Kit (cat. 50671513, Agilent Technologies, Santa Clara, CA, USA) as well as RNA integrity (RIN). Samples with RIN<5.0 were excluded. Reverse transcriptase reaction was performed using SuperScript™ VILO™ cDNA Synthesis Kit (cat. 11754050-ThermoFisher Scientific, Waltham, MA, USA). cDNA deriving from LCM human pancreatic islets was then amplified using TaqMan PreAmp Master Mix (cat. 4488593, ThermoFisher Scientific, Waltham, MA, USA) following manufacturer’s instructions.

Real-Time PCR analysis was performed using TaqMan gene expression assays using primers (see Resources Table) and SensiFast Probe Lo-ROX Kit (cat.# BIO-84020, Bioline) following manufacturer’s recommendation. Data were collected and analysed through Expression Suite software 1.0.1 (ThermoFisher Scientific, Waltham, MA, USA) using 2^-ΔCt^ or 2^-ΔΔCt^ method. ViiA7 Real-Time PCR thermalcycler instrument (ThermoFisher Scientific, Waltham, MA, USA) was used to perform Real-Time PCR reactions.

### 2.8 ACE2 Immunohistochemistry analysis of human pancreatic sections

In order to evaluate the staining pattern of ACE2 in human pancreatic tissues, we analyzed FFPE sections (7-μm thickness), prepared by using a microtome (cat. RM2125 RTS - Leica Microsystems, Wetzlar, Germany) and baked overnight at 37°C, from two different portions of pancreatic tissue for each multiorgan donor (listed in **Table S1**).

After deparaffinization and rehydration through decreasing alcohol series (Xylene-I 20min, Xylene-II 20min, EtOH 100% 5min, EtOH 95% 5min, EtOH 80% 5min, EtOH 75% 5min) pancreatic sections were incubated with 1X Phosphate-Buffered Saline with Ca^2+^ and Mg^2+^ (PBS 1X) supplemented with 3% H2O2 (cat. H1009 - Sigma Aldrich, St. Louis, MO, USA) for 30min, to block endogenous peroxidases. Heat-induced antigen retrieval was performed using 10 mM citrate buffer pH 6.0 in microwave (600W) for 10 minutes, maintaining boiling conditions. Sections were incubated with PBS 1× supplemented with 5% rabbit serum (cat. SCBD33ISV - Sigma Aldrich, St. Louis, MO, USA) to reduce non-specific reactions. Then, sections were incubated overnight at +4°C with primary antibody monoclonal mouse anti-Human ACE2 (cat.MAB933, R&D System, Minneapolis, MN, USA) diluted 1:33 (15 μg/ml) in PBS 1× supplemented with 5% rabbit serum. The next day, sections were incubated with secondary antibody polyclonal rabbit anti-mouse HRP-conjugate (cat.P0260, Dako, Agilent Technologies, Santa Clara, CA, USA) diluted 1:100 in PBS 1× for 1h at room temperature (RT). Subsequently, the sections were incubated with one drop of 3,3’ Diaminobenzidine (DAB) chromogen solution (cat.RE7270-K, Novolink MAX DAB, Leica Microsystems, Wetzlar, Germany) for 5 minutes, to trigger the chromatic reaction. Stained sections were then counterstained with hematoxylin (cat.MHS31, Sigma Aldrich, St. Louis, MO, USA) for 4 minutes, for better visualization of the tissue morphology. After the dehydration through increasing alcohol series, the pancreatic sections were mounted with Eukitt mounting medium (cat.S9-25-37, Bio Optica, Milan, Italy) and covered with a coverslip allowing them to dry. A negative control with only secondary antibody incubation (no primary antibody control sample) was also included in order to exclude potential background artifacts generated by the secondary antibody or the enzymatic detection reaction (**Figure S1A**).

In order to further evaluate ACE2 expression in pancreas sections, the same ACE2 IHC protocol was applied to other 2 primary antibodies anti-Human ACE2: monoclonal rabbit anti-Human ACE2 (cat. Ab108252, Abcam, Cambridge, UK) diluted 1:100 in PBS 1× supplemented with 5% goat serum; and polyclonal rabbit anti-Human ACE2 (cat. Ab15348, Abcam, Cambridge, UK) diluted 1:2000 in PBS 1× supplemented with 5% goat serum. The secondary antibody in both cases was polyclonal goat antirabbit HRP-conjugate (cat. 111-036-003, Jackson ImmunoResearch, Philadelphia, PA, USA), diluted 1:1000 in PBS 1× for 1h at room temperature (RT).

All the 3 primary antibodies anti-Human ACE2, with respective secondary antibodies, were also used to perform a positive control staining in FFPE human lung sections (7-μm thickness), in order to double check the specificity of the primary antibodies.

### 2.9 Immunofluorescence staining for ACE2-Insulin-Glucagon and ACE2-CD31

FFPE pancreatic sections (see above) were analysed by triple immunofluorescence in order to simultaneously evaluate the expression pattern of ACE2, insulin and glucagon. Briefly, after deparaffinization and rehydration through decreasing alcohol series (see above), pancreatic sections were subjected to heat induced antigen retrieval using 10 mM citrate buffer pH 6.0 in microwave (600W) for 10 minutes. Sections were incubated with PBS 1X supplemented with 3% Bovine Serum Albumin (BSA, cat. A1470-25G, Sigma Aldrich, St. Louis, MO, USA) to reduce non-specific reactions. Then, sections were incubated with primary antibody monoclonal mouse anti-human ACE2 (cat. MAB933, R&D System, Minneapolis, MS, USA) diluted 1:33 in PBS 1X supplemented with 3% BSA, overnight at +4°C, followed by polyclonal Rabbit anti-human Glucagon (cat. A0565, Agilent Technologies, Santa Clara, CA, USA) diluted 1:500 in PBS 1X supplemented with 3% BSA, and prediluted polyclonal Guinea Pig anti-human Insulin (cat. IR002 - Agilent Technologies, Santa Clara, CA, USA) as second and third primary antibodies for 1h at room temperature (RT). Subsequently, sections were incubated with goat anti-guinea pig Alexa-Fluor 555 conjugate (cat. A21435, Molecular Probe, ThermoFisher Scientific, Waltham, MA, USA) diluted 1:500 in PBS 1X, goat anti-rabbit Alexa-Fluor 647 conjugate (cat. A21245, Molecular Probe, ThermoFisher Scientific, Waltham, MA, USA) diluted 1:500 in PBS 1X and goat anti-mouse 488 conjugate (cat.A11029 - Molecular Probe, ThermoFisher Scientific, Waltham, MA, USA) diluted 1:500 in PBS 1X, as secondary antibodies for 1h. Sections were counterstained with 4’,6-Diamidino-2-phenylindole dihydrochloride (DAPI, cat. D8517, Sigma-Aldrich) diluted 1:3000 in PBS 1X, and then mounted with Vectashield antifade medium (cat. H-1000, Vector Laboratories, Burlingame, CA, USA) and analysed immediately or stored at +4°C until ready for confocal image analysis. A negative control with only secondary antibody incubation (no MAB933 primary antibody control sample) was also included in order to exclude any background artifacts or fluorochrome overlaps (**Figure S1B**).

For ACE2-CD31 double immunofluorescence staining, FFPE pancreas sections were incubated overnight at +4°C with primary antibody monoclonal mouse anti-human ACE2 (cat. MAB933, R&D System, Minneapolis, MS, USA) diluted 1:33 in PBS 1X supplemented with 3% BSA and with polyclonal rabbit anti-CD31 (cat. Ab28364, Abcam, Cambridge, UK) diluted 1:50 in the same blocking buffer. Subsequently, sections were incubated with goat-anti rabbit Alexa-Fluor 594 conjugate (cat. A11037, Molecular Probe, ThermoFisher Scientific, Waltham, MA, USA) diluted 1:500 in PBS 1X, and goat anti-mouse 488 conjugate (cat. A11029 - Molecular Probe, ThermoFisher Scientific, Waltham, MA, USA) diluted 1:500 in PBS 1X, as secondary antibodies for 1h. Sections were counterstained with DAPI and then mounted as described above.

### 2.10 Cultured cells immunofluorescence

Cultured EndoC-βH1 cells were immunostained for ACE2 and insulin as follows. Cytokines-treated or untreated cells were fixed in 4% PFA for 10 min, washed for 10 min in 0.1 mol/L glycine, permeabilized in 0,25% Triton-X-100 for 5 min and blocked in 3% BSA+0.05% Triton-X100 in PBS without Ca^2+^ and Mg^2+^ for 30 min. EndoC-βH1 cells were incubated with prediluted antibody polyclonal Guinea Pig anti-Human Insulin (cat. IR002 - Agilent Technologies, Santa Clara, CA, USA) for 1h at RT. Then, the cells were washed with PBS without Ca^2+^ and Mg^2+^and incubated with monoclonal mouse anti-Human ACE2 (cat. MAB933, R&D System, Minneapolis, MS, USA) diluted 1:33 in BSA 1% in PBS without Ca^2+^ and Mg^2+^ or with monoclonal rabbit anti-Human ACE2 (cat. ab108252, Abcam, Cambridge, UK) diluted 1:100 in BSA 1% in PBS without Ca^2+^ and Mg^2+^, or with polyclonal rabbit anti-Human ACE2 (cat. ab15348, Abcam, Cambridge, UK) diluted 1:2000 in BSA 1% in PBS without Ca^2+^ and Mg^2+^, and then incubated with primary or with negative isotype control mouse IgG2a (cat. X0943 - Agilent Technologies, Santa Clara, CA, USA) or negative isotype control rabbit IgG (cat. Ab199376, Abcam, Cambridge, UK) for 60 min. Next, EndoC-βH1 cells were rinsed with PBS without Ca^2+^ and Mg^2+^ and incubated with goat anti-mouse-488 1:500 in 1% BSA in PBS without Ca^2+^ and Mg^2+^, or with goat anti-rabbit-488 diluted 1:500 in 1% BSA in PBS without Ca^2+^ and Mg^2+^for 30 min and with goat-anti rabbit Alexa-Fluor 594 conjugate (cat. A11037, Molecular Probe, ThermoFisher Scientific, Waltham, MA, USA). Finally, the cells were incubated with 4’,6-Diamidino-2-phenylindole dihydrochloride (DAPI, cat. D8517, Sigma Aldrich, St. Louis, MO, USA) diluted 1:3000 in PBS 1X and then mounted with Vectashield antifade medium (cat. H-1000 - Vector Laboratories, Burlingame, CA, USA) and analysed immediately or stored at +4°C until ready for confocal image analysis.

HeLa cells were immunostained for ACE2 as follows. The cells were first fixed in 4% PFA for 10 min, washed for 10 min in 0.1 mol/L glycine, permeabilized in 0,25% Triton-X-100 for 5 min and blocked in 3% BSA+0.05% Triton-X-100 in PBS without Ca^2+^ and Mg^2+^ for 30 min. The cells were subsequently washed with PBS without Ca^2+^ and Mg^2+^ and then incubated with monoclonal mouse anti-Human ACE2 (cat. MAB933, R&D System, Minneapolis, MS, USA) diluted 1:33 in BSA 1% in PBS without Ca^2+^ and Mg^2^ or with negative isotype control mouse IgG2a (cat. X0943 - Agilent Technologies, Santa Clara, CA, USA). Then, HeLa cells were rinsed with PBS without Ca^2+^ and Mg^2+^ and were incubated with goat anti-mouse-488 1:500 in 1% BSA in PBS1X without Ca^2+^ and Mg^2+^, or with goat anti-rabbit-488 diluted 1:500 in 1% BSA in PBS1X without Ca^2+^ and Mg^2+^. Cells were counterstained with DAPI and then mounted as described above.

### 2.11 Image analysis

Images were acquired using Leica TCS SP5 confocal laser scanning microscope system (Leica Microsystems, Wetzlar, Germany). Images were acquired as a single stack focal plane or in z-stack mode capturing multiple focal planes (n=40) for each identified islet or selected representative islets. Sections were scanned and images acquired at 40× or 63× magnification. The same confocal microscope setting parameters were applied to all stained sections before image acquisition in order to uniformly collect detected signal related to each channel.

Colocalization analysis between ACE2 and insulin and between ACE2 and glucagon were performed using LasAF software (Leica Microsystems, Wetzlar, Germany). The region of interest (ROI) was drawn to calculate the *colocalization rate* (which indicates the extent of colocalization between two different channels and reported as a percentage) as a ratio between the colocalization area and the image foreground. Evaluation of the signal intensity of ACE2 expression in human pancreatic islets of EUnPOD donors was performed using the LasAf software (www.leica-microsystem.com). This software calculates the ratio between intensity sum ROI (which indicates the sum ROI of the greyscale value of pixels within a region of interest) of ACE2 channel and Area ROI (μm^2^) of human pancreatic islets. Both in colocalization and intensity measurement analysis, a specific threshold was assigned based on the fluorescence background. The same threshold was maintained for all the images in all the cases analysed.

### 2.12 Micro-confocal High-content Screening analysis

Cultured EndoC-βH1 cells were immunostained for ACE2 and insulin as reported above. Cytokines-treated or untreated cells were fixed in 4% PFA for 10 min, washed for 10 min in 0.1 mol/L glycine, permeabilized in 0,25% Triton-X-100 for 5 min and blocked in 3% BSA+0.05% Triton-X100 in PBS without Ca^2+^ and Mg^2+^ for 30 min. EndoC-βH1 cells were incubated with antibody polyclonal Guinea Pig anti-Human Insulin (cat. A21435 - Agilent Technologies, Santa Clara, CA, USA) diluted 1:1740 in BSA 1% in PBS without Ca^2+^ and Mg^2+^and with monoclonal mouse anti-Human ACE2 (cat. MAB933, R&D System, Minneapolis, MS, USA) diluted 1:33 in BSA 1% in PBS without Ca^2+^ and Mg^2+^for 1h at RT or with negative isotype control mouse IgG2a (cat. X0943 - Agilent Technologies, Santa Clara, CA, USA). Then, EndoC-βH1 cells were washed with PBS without Ca^2+^ and Mg^2+^and incubated with goat anti-mouse-488 (cat.A11029 - Molecular Probe, ThermoFisher Scientific, Waltham, MA, USA) and goat anti-guinea pig-555 (cat. A21435, Molecular Probe, ThermoFisher Scientific, Waltham, MA, USA) 1:500 in 1% BSA in PBS without Ca^2+^ and Mg^2+^. EndoC-βH1 cells were incubated with 4’,6-Diamidino-2-phenylindole dihydrochloride (DAPI, cat. D8517, Sigma Aldrich, St. Louis, MO, USA) diluted 1:3000 in PBS 1X; then washed with PBS without Ca^2+^ and Mg^2+^and analysed immediately. Fluorescence images of EndoC-βH1 cells were analysed using Opera Phoenix High Content Screening System (PerkinElmer, Waltham, MA, USA) acquiring multiple images using 63 × magnification; nine microscopic areas per well were automatically selected. Automated image analysis was performed using Harmony^®^ High-Content Imaging (PerkinElmer, Waltham, MA, USA), and fluorescence intensity of treated or untreated cells were measured based on Alexa-555 (insulin) and Alexa-488 (ACE2) fluorochromes. Images were first segmented into nuclei and cytoplasm using the Find Nuclei building block on the DAPI channel and the Find Cytoplasm on the 488 (ACE2) channel. To detect ACE2 signals, Find spots building block was applied to the 488 fluorescence channel inside the cytoplasm area previously detected. The intensity rate was obtained from the average of the nine areas and values reported as Corrected Spot Intensity (which is the “Mean Spot Intensity” minus “Spot Background Intensity”) (31).

### 2.13 RNA sequencing processing and analysis

Total RNA of EndoC-βH1 cells and of pancreatic human islets exposed or not to IFNα or to IL-1β + IFNγ for the indicated time points was obtained and prepared for RNA sequencing as described (32–34). Bioanalyzer System 2100 (Agilent Technologies, Wokingham, UK) was used to evaluate samples quality by determining RNA integrity number (RIN) values. Only samples presenting RIN values > 9 were analyzed. The obtained libraries were submitted to a second quality control before sequencing on an Illumina HiSeq 2500. The Salmon software version 0.13.2 (35) was used to re-analyse our original RNA-seq data (32–34) by mapping the sequenced reads to the human reference transcriptome from GENCODE version 31 (GRCh38) (36) using the quasi-alignment model. Gene expression is represented in Transcripts Per Million (TPM).

Differentially expressed genes were identified with DESeq2 version 1.24.0 (37). The estimated number of reads obtained from Salmon were used to run the DESeq2 pipeline. In summary, in this approach the DESeq2 normalizes samples based on per-sample sequencing depth and accounting for the presence of intra-sample variability. Next, data were fit into a negative binomial generalized linear model (GLM) and computed using the Wald statistic. Finally, obtained p-values were adjusted for multiple comparisons using the false discovery rate (FDR) by the Benjamini-Hochberg method (38). Genes were considered significantly modified with a FDR < 0.05.

### 2.14 Western blot analysis

Total proteins from EndoC-βH1 cells were extracted using a lysis buffer (20 mM Tris-HCl pH 8.0, 137 mM NaCl, 1% (w/v) NP-40, 2mM EDTA) supplemented with 1X protease inhibitors (Roche). Total proteins were quantified using Bradford assay, and 50-100 μg protein/lane were separated using SDS-PAGE Tris-Glycine gradient Bis-Acrylamide gel 4-20%. Proteins were then transferred to Nitrocellulose 0.2 μm membrane using wet electrophoresis system. Upon transfer onto nitrocellulose, membranes were washed 3 times with TBST1X (Tris-HCl 25 mM, NaCl 150 mM, Tween 20 0,1%, pH 7.4) and then incubated 2h with 5% non-fat dry milk in TBST1X. To identify ACE2, three different antibodies were used: #Ab108252, #Ab15348 (Abcam) and #MAB933 (R&D system) were respectively diluted 1:1000, 1:500 and 1:250 in 5% non-fat dry milk in TBST1X and incubated o/n at +4°C and then with Goat anti-rabbit (#111-036-003, Jackson Laboratories) or Goat anti-Mouse (#115-036-003, Jackson Laboratories) diluted 1:5000 in 2% non-fat dry milk in TBST1X 1 h RT. After 3 washes with TBST1X and 1 wash in TBS1X, chemiluminescent signal was detected by using ECL solution (GE Healthcare, Little Chalfont, Buckinghamshire, UK-RPN2232). Chemiluminescent analysis of immunoblot results was performed by using LAS400 analyzer (GE Healthcare, Little Chalfont, Buckinghamshire, UK-RPN2232).

### 2.15 ACE2 targeted Mass Spectrometric (MS)-Shotgun proteomics analysis

To perform ACE2 targeted MS analysis, EndoC-βH1 cells were lysed with RIPA buffer 1X and protein lysate concentration quantified through BCA assay. Then the protein lysate was mixed with 400 μL of urea 8M in Tris-HCl 100nM pH 8,5 (UA), with the addition of 100 mM DTT. The mixture was charged on a filter 10 K Pall, incubated 30min RT and centrifuged 13,800xg 30min. The filter was washed twice with 400 μL of UA and centrifuged 13,800xg 30min, then incubated with 100 μL of 50 mM of iodoacetamide (IAC) solution in a thermo-mixer for 1min 150 RPM and without mixing for 20min, then centrifuged at 13,800xg for 20min. After these steps, the filter was washed twice with 400 μL of UA and centrifuged at 13,800xg 30min, twice with 400 μL of 50 mM ammonium bicarbonate (AMBIC), and then centrifuged twice, a first time at 13,800xg for a30min and a second time at 13,800xg for 20min. Next, 40 μL of 50 mM AMBIC were added to the filter together with trypsin (ratio trypsin/proteins 1:25) and incubated O/N 37°C. The sample was then transferred into a new collecting tube and centrifuged at 13,800xg for 10min. Subsequently, 100 μL of 0.1% formic acid was added on the filter and centrifuged 13,800xg 10min. Finally, filter the was discarded and the solution was desalted with OASIS cartridges according to manufacturers’ instructions. The retrieved peptides were concentrated through SpeedVac and the sample was resuspended in a solution of 3% acetonitrile, 96.9% H2O and 0.1% formic acid. The analyses were performed on a Q-Exactive Plus mass spectrometer (Thermofisher Scientific), equipped with electrospray (ESI) ion source operating in positive ion mode. The instrument is coupled to an UHPLC Ultimate 3000 (Thermofisher Scientific). The chromatographic analysis was performed on a column Acquity UPLC Waters CSH C18 130Å (1 mm X 100 mm, 1,7 μm, Waters) using a linear gradient and the eluents were 0.1% formic acid in water (phase A) and 0.1% formic acid in acetonitrile (phase B). The flow rate was maintained at 100 μl/min and column oven temperature at 50°C. The mass spectra were recorded in the mass to charge (m/z) range 200-2000 at resolution 35K at m/z 200. The mass spectra were acquired using a “data dependent scan”, able to acquire both the full mass spectra in high resolution and to “isolate and fragment” the ten ions with highest intensity present in the full mass spectrum. The raw data obtained were analyzed using the Biopharma Finder 2.1 software from ThermoFisher Scientific.

The elaboration process consisted in the comparison between the peak list obtained “in silico” considering the expected aminoacidic sequence of human ACE2 protein (Uniprot ID: Q9BYF1), trypsin as digestion enzyme and eventual modifications (carbamidomethylation, oxidation, etc.).

### 2.16 ACE2 promoter transcription factors (TF) binding motifs analysis

ACE2 proximal promoter sequence was retrieved from Ensembl Genome browser database (http://www.ensembl.org/index.html) [Release 100 (April 2020)]. ACE2 gene (ENSG00000130234) was searched in Human genome GCRh38.p13 assembly. The sequence of interest was retrieved using “Export Data” function by selecting 1000 bp upstream 5’ Flanking Sequence (GRCh38:X:15602149:15603148:-1) and downloaded in FASTA format. The analysis of TF binding motifs was performed using Transcription factor Affinity Prediction (TRAP) Web Tool (http://trap.molgen.mpg.de/cgi-bin/trap_receiver.cgi)(39). In TRAP, ACE2 promoter sequence was analysed by using both TRANSFAC.2010.1 and JASPAR vertebrate databases and *human_promoters* as background model.

### 2.17 Data and Code Availability

The NCBI GEO accession number for RNA sequencing data reported in this paper are: GSE133221, GSE108413, GSE137136.

### 2.18 Statistical analysis

Results were expressed as mean ± SD. Statistical analyses were performed using Graph Pad Prism 8 software. Comparisons between two groups were carried out using Mann-Whitney U test (for non-parametric data) or Wilcoxon matched-pairs signed rank test. Differences were considered significant with p values less than 0.05.

## 3 Results

### 3.1 ACE2 expression pattern in human pancreas

To determine the ACE2 protein expression pattern in human pancreatic tissue, we first performed a colorimetric immunohistochemistry analysis to detect ACE2 on formalin-fixed paraffin embedded (FFPE) pancreatic sections obtained from seven (n=7) adult non-diabetic multiorgan donors collected by the INNODIA EUnPOD biobank (**Table S1**). To specifically detect ACE2 protein in such context, we initially used a previously validated monoclonal anti-human ACE2 antibody (R&D MAB933) (7) which passed the validation criteria suggested by the International Working Group for Antibody Validation (IWGAV) (7,40) (see ***Resources Table* in Supplementary Material**). For each pancreas, two sections derived from two different FFPE tissue blocks belonging to different parts of the organ (head, body or tail) were analysed. Based on pancreas morphometry and histological composition, we identified three main cell types positive for ACE2 (**Figure 1, panel-a to -f**). In the exocrine pancreas there was a marked and intense staining in a subset of vascular components (endothelial cells or pericytes) found in inter-acini septa (**Figure 1, panel-a and -b**). We also identified ACE2 positive cells in the pancreatic ducts, even though only some scattered cells with a clear ACE2 signal were detected (**Figure 1, panel-c and -d**). Of interest, we observed a peculiar ACE2 staining pattern in the endocrine pancreatic islets, showing a diffuse ACE2 signal in a subset of cells within islet parenchyma (**Figure 1, panel -e and -f; Figure S2**). However, the observed ACE2 expression in islets was lower than the expression observed in microvasculature, the latter representing the main site for ACE2 expression in the pancreas.

**Figure 1.**
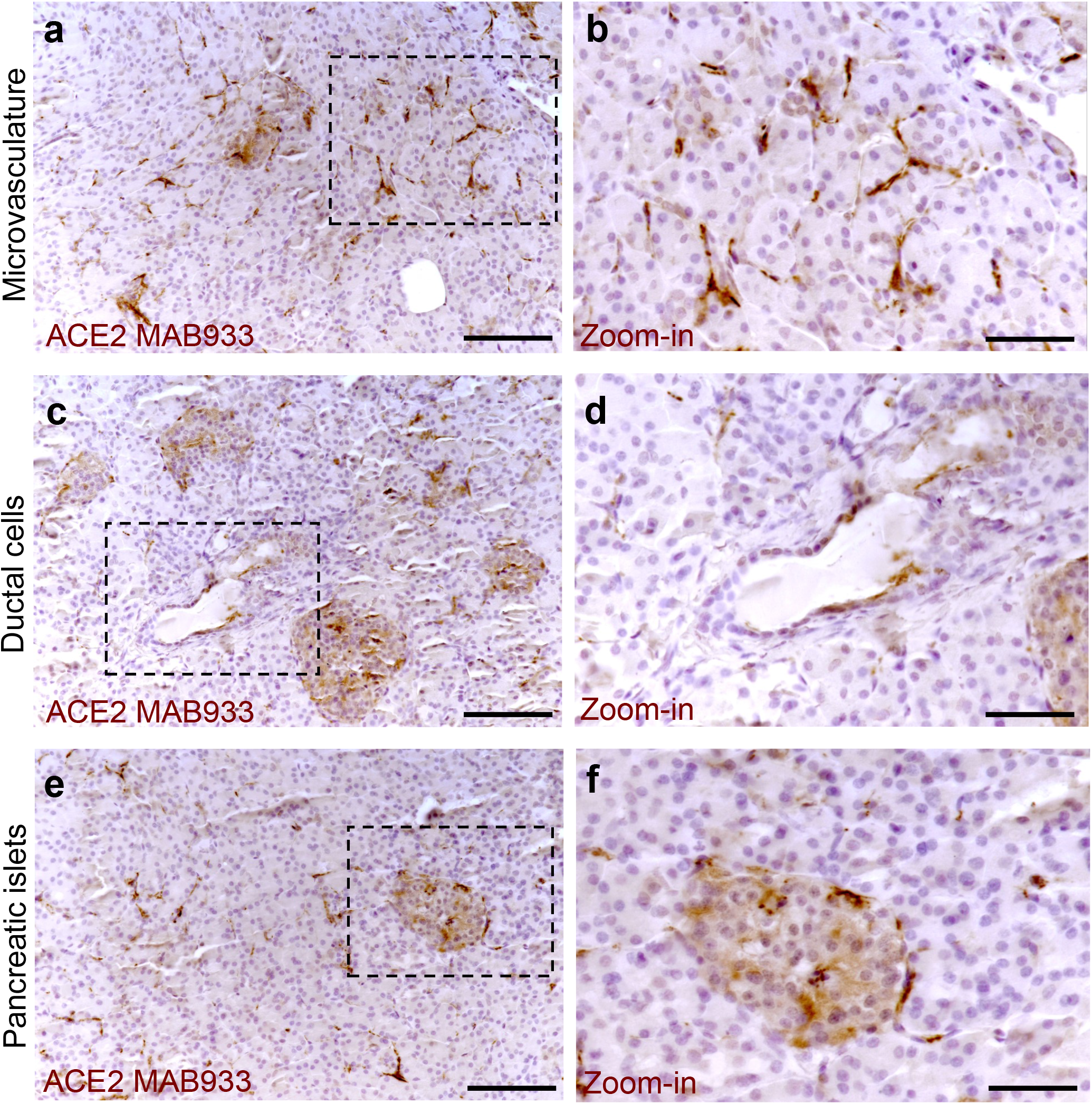
ACE2 staining pattern in human pancreas. Immunohistochemistry for ACE2 in human pancreatic tissue sections (case #110118) using R&D MAB933 antibody. ACE2 is markedly expressed in microvasculature associated cells (panel-a and -b), in some rare ductal cells (panel-c and -d) and in a subset of endocrine cells within pancreatic islets (panel-e and -f). Scale bars in panel-a, -c and -e: 150 μm. Scale bars in panel-b, -d and-f: 70 μm. Zoom-in images are reported in panel -b, -d and -f

In all cases analysed, including different blocks of the same case, a similar expression pattern of ACE2 was observed, even though a certain degree of variability in terms of ACE2 staining intensity within the islets was noted (**Figure S2**).

The highest signal of ACE2 within the pancreas was observed in putative association with microvasculature (**Figure 1, panel-a and -b**). Of note, in such context, a lobular staining pattern of microvasculature associated ACE2 was evident, as demonstrated by the presence of positive cells in certain lobules and low or null expression in other lobules of the same pancreas section (**Figure 2A**). ACE2 staining pattern in inter-acini septa suggested an overlap with cells associated to microvasculature, most likely endothelial cells. In order to explore such possibility, we performed a double immunofluorescent staining on pancreas FFPE sections for ACE2 and the endothelial cell specific marker CD31. The results showed that ACE2 signal is associated, but not superimposed, to the CD31-specific one, thus resembling the tight association of pericytes to endothelial cells and strongly suggesting the presence of ACE2 in microvasculature pericytes (**Figure 2B**).

**Figure 2.**
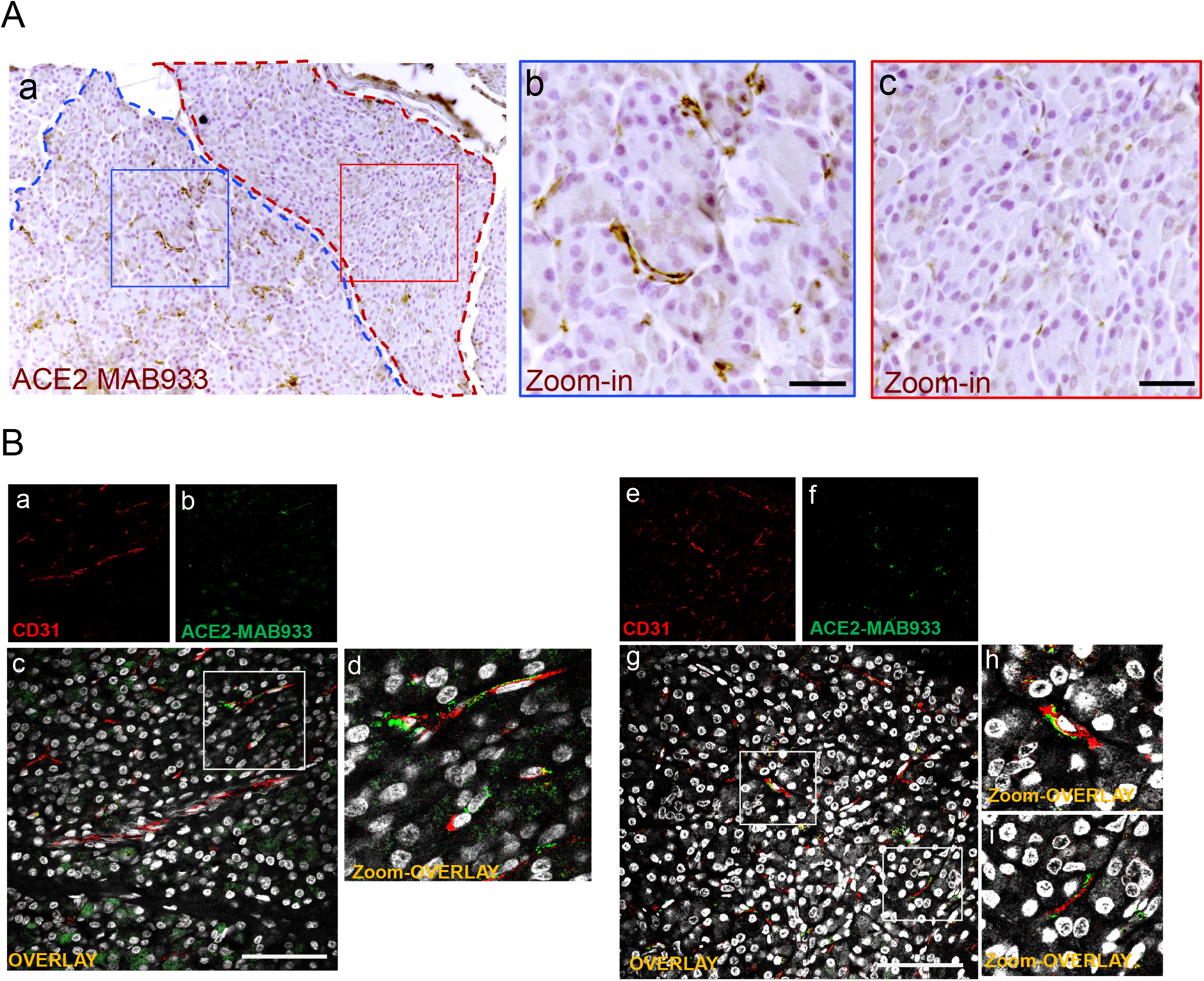
In microvasculature, ACE2 is putatively expressed in pericytes. (**A**) Representative image of human pancreatic Formalin-Fixed Paraffin Embedded (FFPE) section stained for ACE2 in case #301118. In panel-a, a representative image of a pancreatic section showing two adjacent lobules (blue and red dotted lines) with different staining for ACE2 in endothelial cells/pericytes. A specific segmentation of the two lobules with high (blue) (zoom-in, panel-b) and low or null expression of ACE2 (red) (zoom-in, panel-c) is shown, suggesting lobularity of ACE2 expression in exocrine endothelial cells/pericytes of human pancreas. Scale bar in panel-a: 100 μm. Scale bar in panel-b and -c: 30 μm. (**B**) Double immunofluorescence staining of ACE2 (green) and CD31 (red) in FFPE pancreas sections from Body01A of Case #110118 (panel-a to −d) and of Body01B of Case #141117 (panel-e to −i). Digital zoom-in overlay images are shown in panels -d, -h and −i. Scale bar in panel-d and −g: 100 μm.

In order to confirm the ACE2 cellular distribution observed in the pancreas using MAB933 antibody, we tested two additional anti-ACE2 antibodies from Abcam: Ab15348 and Ab108252 (see ***Resources Table***). According to the informations obtained by R&D and Abcam, while MAB933 and Ab15348 are reported to recognize the C-terminus portion of ACE2 (18-740aa and 788-805aa, respectively), Ab108252 is specifically designed to react with a linear peptide located in the N-terminal ACE2 protein sequence (200-300aa). It has been recently reported that a 459 aa short-ACE2 isoform (357-805aa of ACE2 + 10aa at N-terminus) can be co-expressed alongside the full-length ACE2 protein (1-805aa) (**Figure 3A**) (24,25); the short-ACE2 misses part of the N-terminal region targeted by Ab108252 antibody. We observed ACE2 islet-related signal using both MAB933 (**Figure 3B**) and Ab15348 (**Figure 3C**), but the Ab108252 antibody did not show any positivity within the islet parenchyma (**Figure 3D**), raising the possibility that the most prevalent ACE2 isoform within pancreatic islets is the short one. Of note, the three antibodies tested showed ACE2 positivity in the microvasculature, thus suggesting a putative differential distribution of the two ACE2-isoforms in the human pancreas.

**Figure 3.**
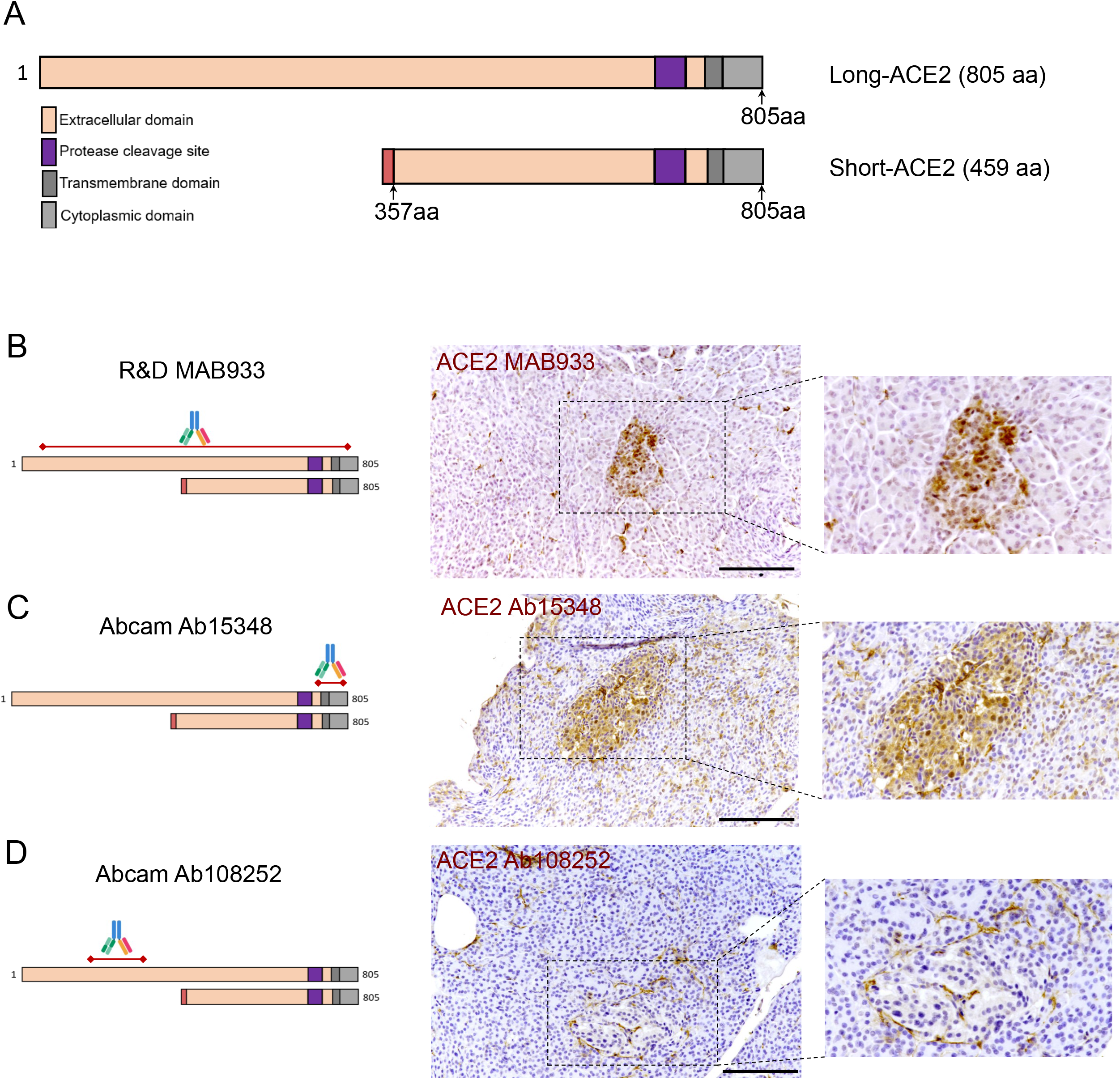
ACE2 immunohistochemistry staining pattern in human pancreatic islets using MAB933, Ab15348 and Ab108252 antibodies. (**A**) Aligned sequences and structures of recently described ACE2 isoforms, long-ACE2 (805aa, ~110 kDa) and short-ACE2 (459aa, ~50kDa). Main ACE2 protein domains are reported with different colours. (**B**) R&D MAB933, (**C**) Abcam Ab15348 and (**D**) Ab108252 antibody predicted target sequence within the two ACE2 isoforms, alongside with immunohistochemistry staining distribution in pancreatic islets. Scale bars in (B), (C) and (D) are 100μm.

As a positive control for our immunohistochemistry method and ACE2 antibodies adopted, we evaluated FFPE lung tissue sections. As previously shown (7,41,42), we observed scattered positive cells (putatively AT2 pneumocytes) in the alveolar epithelium both using MAB933 and Ab 15348 (**Figure S3A** and **S3B**). In contrast, we did not observe any signal using Ab108252 (**Figure S3C**). Collectively, these results indicate that the same staining pattern were obtained by using 2 out of 3 antibodies that may recognize both ACE2 isoform (short-ACE2 and long-ACE2) thus confirming: *(i)* a high ACE2 expression in microvasculature pericytes; (ii) rare scattered ACE2 positive ductal cells; (*iii*) diffuse though weak ACE2 positive staining in a subset of cells within human pancreatic islets.

### 3.2 In human pancreatic islets ACE2 is preferentially expressed in β-cells

Using both MAB933 and Ab15348 we observed ACE2 signal in pancreatic islets which suggests that ACE2 is expressed in endocrine cells. Therefore, we sought to determine which pancreatic islet cell subset contributes to ACE2 signal in such context. To do so, we performed a triple immunofluorescence analysis on the same set of FFPE pancreatic sections of non-diabetic multiorgan donors, aimed at detecting glucagon-positive α-cells, insulin-positive β-cells and ACE2 signals (**Figure 4 and Figure S4**).

**Figure 4.**
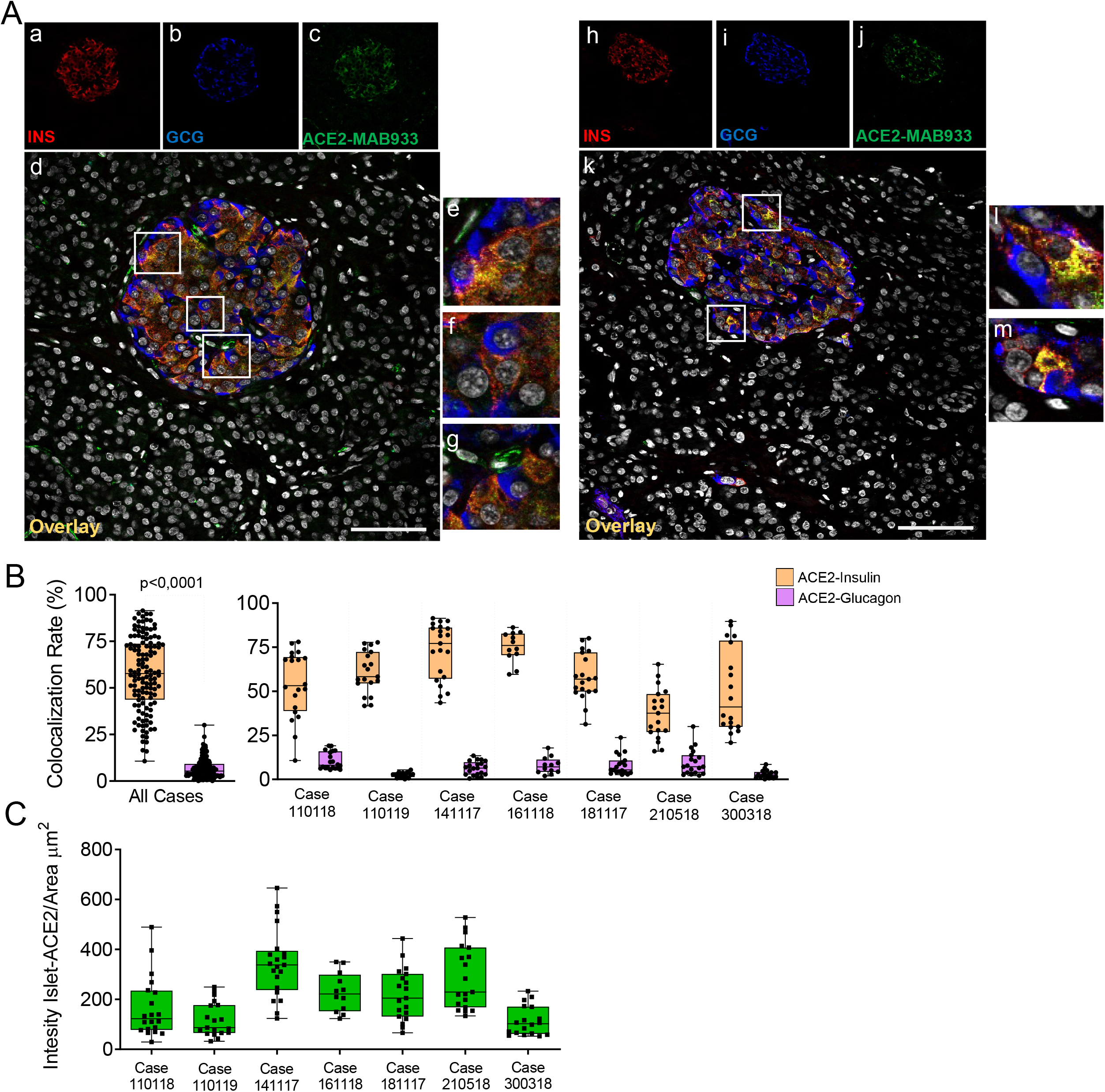
In human pancreatic islets, ACE2 is preferentially expressed in insulin-producing β-cells. Triple immunofluorescence staining and image analysis of FFPE human pancreatic section stained for insulin (red), glucagon (blue) and ACE2 (green). (**A**) Representative islets of two different cases. Panel-a to -g: representative pancreatic islet of FFPE pancreas block Body01A of case #110118. Panel-h to -m: representative pancreatic islet of FFPE pancreas block Body01B of case #141117. Panel-e to -g: digital zoom in images of the pancreatic islet shown in panel-d. Panel-l and -m: digital zoom in images of the pancreatic islet shown in panel-k. Scale bar in panel-d and -k: 100 μm. (**B**) Colocalization rate analysis of overlapping ACE2-insulin and ACE2-glucagon in 128 single pancreatic islets of 7 different cases. p-value was calculated using Wilcoxon matched-pairs signed rank test. On the right: colocalization rate analysis of ACE2-insulin and ACE2-glucagon in each of the 7 cases analysed. For each case, a total of 7-11 islets/section were analysed. (**C**) Analysis of the intensity of ACE2 islet-related signals in the cases analyzed. Values are shown as fluorescence intensity of each islet reported as the sum of gray-scale values for each pixel normalized for the islets area (ROI, μm^2^).

Using R&D MAB933, ACE2 preferentially overlapped with the insulin-positive β-cells (**Figure 4A, panel-a to -m**), being mostly colocalized with insulin and low/not detectable in α-cells (**Figure 4A, panel-e, -f, -l, -m**). Such staining pattern was observed in all cases and was consistent between two different FFPE pancreas blocks of the same case (**Table S2**). As expected, ACE2-only positive cells within or around pancreatic islets were also observed (**Figure 4A, panel g**), potentially indicating the presence of ACE2-positive pericytes interspersed in the islet parenchyma or surrounding it.

Intriguingly, in the β-cells a major fraction of ACE2 was observed in the cytoplasm and partially overlapped with insulin positive signal, while in a subsets of them only a minor fraction of the ACE2 signal was attributable to several spots located on plasma membrane (**Figure 4A; Figure S5A red arrow, and Figure S5B**). In microvasculature pericytes, the ACE2-positive signal was mainly observed in plasma membrane (**Figure 4A, panel g; Figure S5 green arrow)** as previously described (43). There were also some ACE2-negative β-cells (**Figure S5, white arrow**).

Colocalization rate analysis between ACE2-insulin and ACE2-glucagon, performed on a total of 128 single pancreatic islets from seven different adult non-diabetic cases, confirmed the significant preferential expression of ACE2 in β-cells compared to α-cells (colocalization rate: ACE2-INS 57.6±19.3% vs. ACE2-GCG 6.8±5.4 % p<0.±0001) (**Figure 4B and Figure S6A**) The comparison of colocalization rates between ACE2-insulin and ACE2-glucagon among all cases analysed, confirmed the consistent preferential expression of ACE2 in β-cells in comparison to α-cells (**Figure 4B**). These results were confirmed when comparing different blocks of the same case (**Table S2**). There was however heterogeneity in terms of the ACE2-insulin colocalization rate among different islets (ACE2-INS colocalization rate range: 0.6 - 91.4%). Such heterogeneity was also highlighted by the presence of rare ACE2-negative pancreatic islets in the same pancreas section.

Inter-islets heterogeneity was also clearly observed regarding ACE2 islet-related signal intensity analysis (**Figure 4D**).

Of note, some cases showed a lower ACE2-insulin mean colocalization rate and islet-ACE2 signal intensity compared to the other ones (**Figure 4D**), thus suggesting a high degree of heterogeneity among cases also in terms of islet-ACE2 expression. No significant correlation between ACE2-islets signal intensity and Age, BMI or cold-ischemia time were observed in our donors cohort (**Figure S6B**).

### 3.3 A short ACE2 isoform is prevalently expressed in the human β-cell line EndoC-βH1

Using Western Blot (WB) and immunofluorescence analysis, we explored the expression of ACE2 in the human β-cell line EndoC-βH1, a model of functional β-cells for diabetes research (27,44). To do so we used R&D MAB933, Abcam Ab15348 and Ab108252 antibodies, as previously done in the above described pancreas immunohistochemistry experiments. In WB analysis, MAB933 revealed the presence of a prevalent 50 kDa band corresponding to the short-ACE2 isoform; use of this ab showed the brightest signal in immunofluorescence staining among the three antibodies tested (**Figure 5A**). Abcam Ab15348 worked better in WB for the recognition of both ACE2 isoforms, and indicated that the most prevalent ACE2 isoform present in human β-cells is the short-ACE2 (50 kDa, blue arrow) (**Figure 5B**). In contrast, Ab108252 recognized only the long-ACE2 isoform (>110 kDa-red arrow) (**Figure 5C**). Of note, the results obtained through WB analysis are in line with the immunofluorescence signal which revealed that Ab108252 only stained a minor fraction of EndoC-βH1 and the obtained signal was mainly found on the plasma membrane (**Figure 5C, panel-b**). Conversely, Ab15348 and MAB933, which recognized both ACE2 isoforms, showed a higher signal and a different subcellular localization respect to Ab108252 (**Figure 5A, 5B, panel-b**). MAB933 ACE2-insulin double immunofluorescence staining confirmed the main punctuate and likely granular cytoplasmic ACE2 signal which also partially overlapped with insulin-positive secretory granules (**Figure 5D, panel -a to panel -h**). In addition, we also observed some spots putatively localised on the plasma membrane (**Figure 5D**). Of note, the specificity of ACE2 MAB933 signal observed in EndoC-βH1 was orthogonally tested in comparison to HeLa cells which showed very low/absent ACE2 mRNA expression (**Figure S7A**) and resulted indeed negative for ACE2 in immunofluorescence (**Figure S7B**).

**Figure 5.**
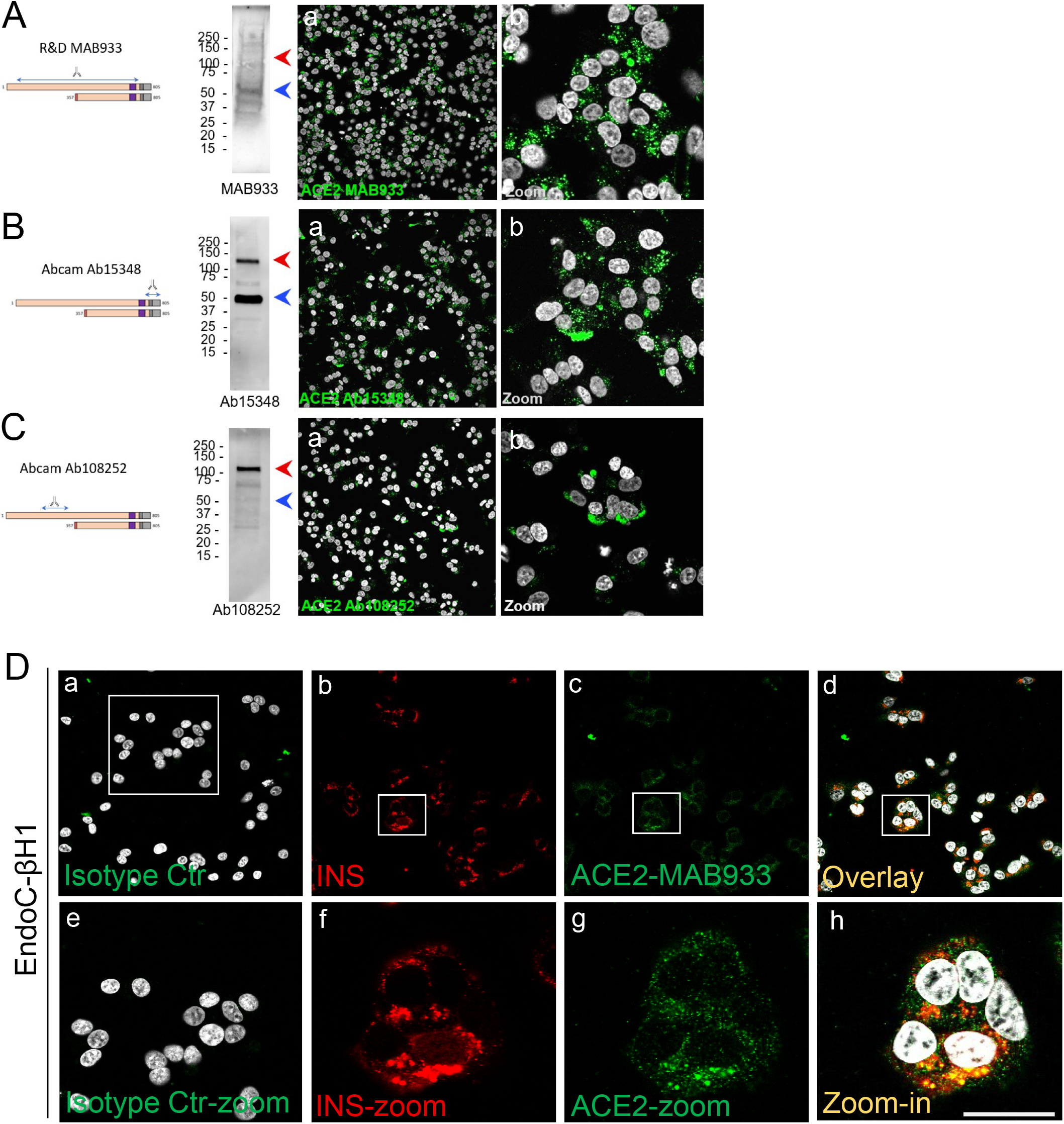
A short-ACE2 isoform is prevalently expressed in human β-cell line EndoC-βH1. Western Blot and Immunofluorescence analysis of EndoC-βH1 using (**A**) R&D monoclonal MAB933, (**B**) Abcam polyclonal Ab15348 and (**C**) Abcam monoclonal Ab108252 anti-ACE2 antibody. For each antibody adopted, specific target sequence is reported within the aligned ACE2 isoforms. In western blot analysis, molecular weight markers (from 15 kDa to 250 kDa) are reported; red arrows indicate long-ACE2 isoform (expected band of ~110kDa), while blue arrows indicate short-ACE2 isoform (~50kDa). (**D**) ACE2 (R&D MAB933) and insulin double immunofluorescence analysis in EndoC-βH1 cultured cells. Negative isotype primary antibody control (relative to ACE2 primary antibody) is shown in panel-a. Insulin (red) and ACE2 (green) are reported in panel-b and -c, while overlay is reported in panel-d. Digital zoom-in images are reported from panel-e to -h. Scale bar in panel -h = 15 μm.

An additional evidence of the presence of ACE2 in EndoC-βH1 was provided by the shotgun proteomic analysis, aimed at detecting specific peptides derived from ACE2 protein independently of the use of specific antibodies. By this independent approach, we observed the presence of both N-terminal and C-terminal unique ACE2-derived peptides (**Figure S8**), which further confirmed the presence of the ACE-2 protein in human β-cells (**Supplementary File 1a, 1b**).

### 3.3 Total ACE2 mRNA is expressed in human pancreatic islets and in the human beta-cell line EndoC-βH1

To confirm the ACE2 expression in human islets, we also evaluated its transcriptional activity both in collagenase-isolated and in Laser-Capture Microdissected (LCM) human pancreatic islets, by measuring its mRNA expression using TaqMan RT-Real Time PCR. In order to avoid detection of genomic DNA, we used specific primers set generating an amplicon spanning the exons 17-18 junction of ACE2 gene, thus uniquely identifying its mRNA (**Figure S9A**). Of note, the selected amplicon is shared between short and long-ACE2 isoforms thus identifying total ACE2 mRNA.

First, as a positive control we analysed total ACE2 expression in RNA extracted from a lung parenchyma biopsy tissue (**Figure 6A and 6E**). Collagenase-isolated human pancreatic islets obtained from four different non-diabetic donors pancreata (**Table S1**) showed ACE2 mRNA expression, as demonstrated by RT-Real-Time PCR raw cycle threshold (Ct) values, reporting a Ct range between 28-29 (**Figure 6B and 6F**). Since human pancreatic islets enzymatic isolation procedures may induce some changes in gene expression (45), we microdissected human islets from frozen pancreatic tissues obtained from five non-diabetic multiorgan donors recruited within INNODIA EUnPOD network (46) and evaluated ACE2 mRNA levels. The LCM procedure (**Figure S9B**) allowed us to extract high quality total RNA (**Figure S9C**) from human pancreatic islets directly obtained from their native microenvironment, thus maintaining transcriptional architecture. ACE2 mRNA expression in LCM-human pancreatic islets showed a consistent expression among cases, similar to isolated islets, as shown by ACE2 mRNA raw Ct and normalized values (**Figure 5C and 5G**).

**Figure 6.**
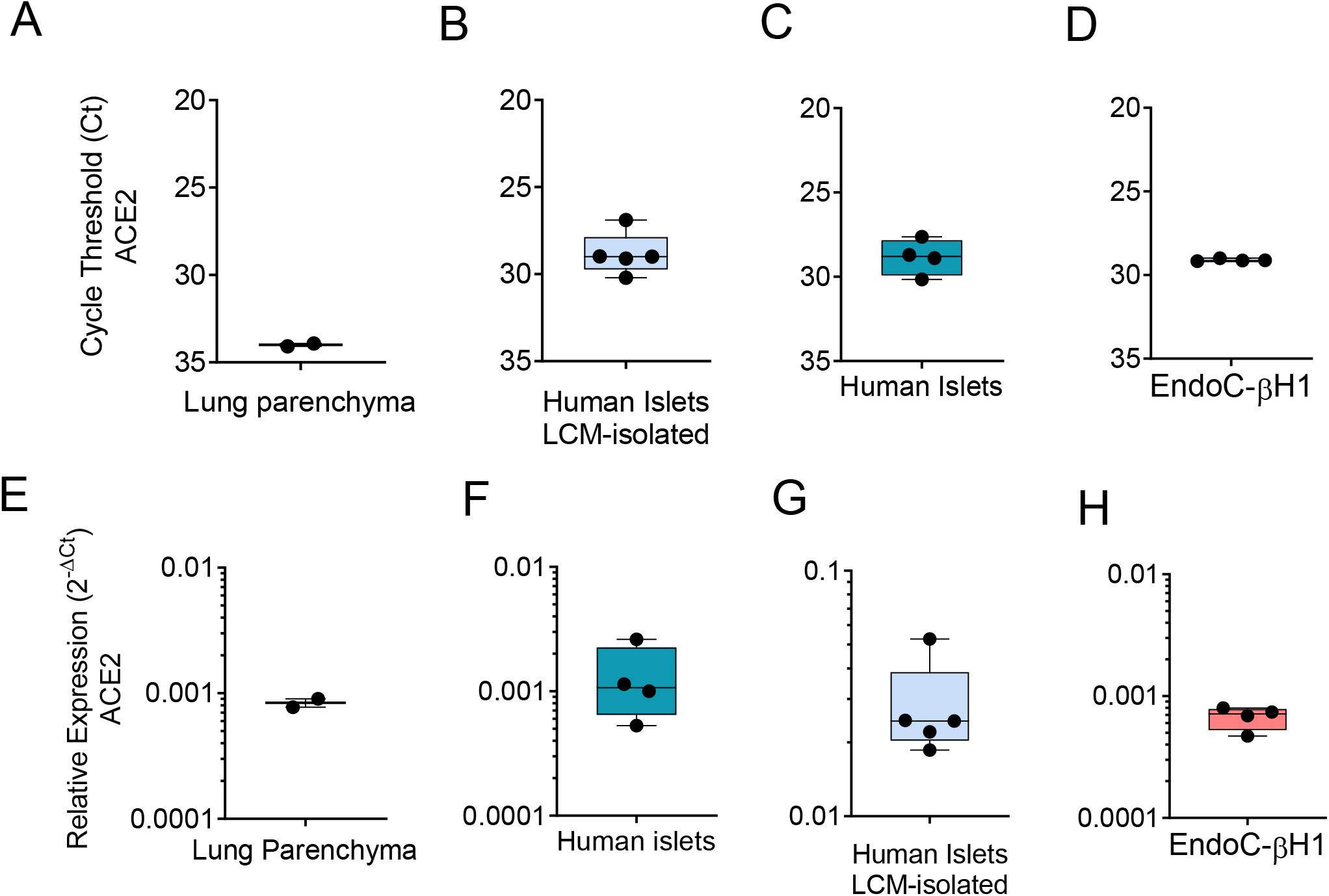
qRT-Real Time PCR analysis of ACE2 mRNA. (**A-D**) ACE2 raw Ct values results in lung tissue (n=1, in duplicate), in enzymatic-isolated human pancreatic islets samples (n=4), in LCM-microdissected islets (n=5) and in EndoC-βH1 (n=4). (**E-G**) ACE2 expression values normalized using GAPDH and β2-microglobulin of the samples analysed in A-D. Values are reported as 2^-dCt^. Mean ± S.D. values are shown.

**Figure 7.**
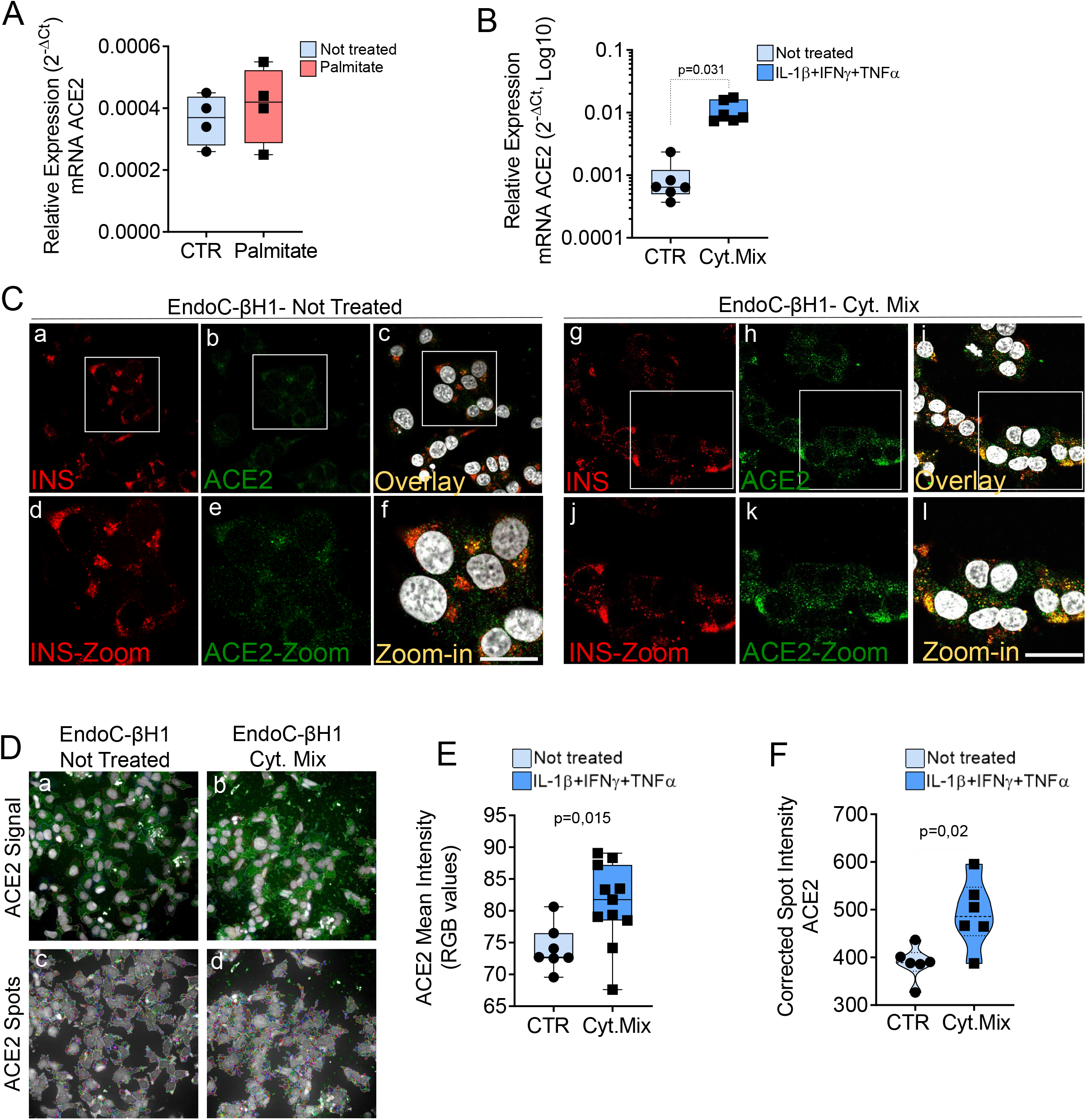
ACE2 expression is increased by inflammatory stress. ACE2 mRNA RT-Real-Time PCR analysis in EndoC-βH1 treated or not (CTR) with Palmitate (2,0 mM) (**A**) or with cytokines (IL-1β+IFNγ+TNFα) (Cyt. Mix) (**B**) for 24h. Results are reported as mean ± S.D of 2^-dCT^ normalized values. p-values were calculated using Wilcoxon matched-pairs signed rank test. (**C**) Immunofluorescence analysis of insulin (red, panel-a and -g) and ACE2 (green, panel-b and -h) in EndoC-βH1 not-treated (panel-a to -f) or treated (panel-g to -l) with cytokines for 24 h. Digital zoomin images are reported in panel-d to -f and in panel-j to -l. Scale bar in panel-f: 10 μm. Scale bar in panel-l: 15 μm. (**D**) Representative images of ACE2 staining (green) and analysis using micro-confocal High-content screening in EndoC-βH1 treated or not with cytokines (IL-1 β+IFNγ+TNFα). Panel-a and -b: ACE2 signal (green) and automated cell cytoplasm segmentation and identification. Panel -c and -d: identification of ACE2 granular spots within segmented cytoplasm in panel-a and -b. Each ACE2 granular spot intensity was measured and analysed. (**E**) Mean intensity imaging analysis related to data reported in (C) of EndoC-βH1 treated or not with cytokines. Data are reported as individual values alongside with mean ± S.D of RGB grey-intensity measures of 6-11 different experimental points related to two different independent experiments. Individual values alongside with mean ± S.D are reported. P-value was calculated using Mann Whitney U test. (**F**) High content screening analysis of Corrected Median Spot intensity of ACE2 signal in EndoC-βH1 treated or not with cytokines. Median intensity values of 6 different experimental points are reported. P-value was calculated using Mann Whitney U test (p<0,05).

Finally, we analysed total ACE2 mRNA expression in the human β-cell line EndoC-βH1. Analysis of ACE2 mRNA expression in these cells demonstrated a similar expression level in comparison to human pancreatic islets (**Figure 3D and 3H**), with raw Ct values ranging from 28 to 30.

### 3.4 ACE2 expression is increased in EndoC-βH1 cells and in human pancreatic islets by pro-inflammatory cytokines

In order to determine whether metabolic or inflammatory stress conditions modify pancreatic endocrine β-cell expression of ACE2, we exposed the human β-cell line EndoC-βH1 and isolated human pancreatic islets to metabolic or inflammatory stressors and subsequently evaluated ACE2 expression levels.

Exposure to palmitate (2mM palmitate for 24 h) did not significantly modulate ACE2 expression (**Figure 6A**). In line with these observations, neither primary human islets exposed to palmitate (47) nor human islets isolated from patients affected by type 2 diabetes (48) and evaluated by RNA sequencing showed any increase in ACE2 mRNA expression as compared to the respective controls (respectively 0.9 control vs 0.5 palmitate and 2.4 control vs 3.5 T2D; data reported as Trimmed Means of M values (TMM); not significant). On the other hand, upon 24 h exposure to a pro-inflammatory cytokines mix (IL-1β-IFNγ and TNFα), EndoC-βH1 cells significantly upregulated ACE2 mRNA levels (fold change: 12.3 vs. not-treated control, p=0.031) (**Figure 6A**). The same results were confirmed through immunofluorescence analysis aimed at measuring ACE2 protein levels and subcellular localization in EndoC-βH1 exposed or not to the same proinflammatory condition (**Figure 6C**). Indeed, we observed a significant increase in ACE2 mean intensity values upon cytokine treatment, confirming the upregulation of ACE2 protein as well (**Figure 6E**). These results were confirmed using an automated micro-confocal high content images screening system (31) which allowed us to measure ACE2 intensity in cytokine-treated vs not-treated EndoC-βH1 cells (**Figure 6D** and **Figure 6F**).

In support of the observed increase of ACE2 upon pro-inflammatory stress, RNA sequencing data analysis of EndoC-βH1 cells exposed to IL-1β+IFNγ (48h) or to IFNα (18h) further confirmed such increase (**Table S3**). Indeed, we observed a 24.5 and 55.2 fold-increase (p<0.0001) in total ACE2 mRNA [transcript (ENST00000252519.8)] (**Table S3 and Figure S10A**) expression in EndoC-βH1 cells treated with IL-1β+IFNγ or with IFNα, respectively. Importantly, the same expression pattern was observed also in human pancreatic islets exposed to the same cytokines mix, as demonstrated by a 2.4 and 5.1 fold-increase in ACE2 mRNA expression following IL-1β+IFNγ or IFNα treatment respectively (p<0.0001) (**Table S3 and Figure S10B**).

Additionally, in order to strengthen such observations, we focussed on the ACE2 gene promoter by analysing its upstream sequence (−1000 bp), from ACE2 transcriptional start site (TSS). Using two different transcription factors (TF) binding motifs databases, we found several binding sites for TFs related to cytokine signalling pathways such as STAT1 or STAT3 (**Figure S11**), thus reinforcing our results of an association between inflammation and ACE2 expression, and confirming what previously reported (65). However, the analysis of ACE2 expression distribution in FFPE pancreas sections from a T1D longstanding donor (see **Table S1**) did not show remarkable changes in the levels or distribution of ACE2 in infiltrated islets (**Figure S12**). This is in line with RNAseq analysis of whole islets from two T1D patients and four controls, which indicated similar ACE2 expression (RPKM 1.2-1.7 in all cases) (49). Analysis of additional recent-onset T1D donors is needed to evaluate potential changes in ACE2 expression in β-cells of highly infiltrated pancreatic islets.

Collectively these results demonstrate that ACE2 is upregulated upon *in vitro* exposure to early and acute inflammatory, but not metabolic, stressors both in EndoC-βH1 and in human pancreatic islets.

## 4 Discussion

In COVID-19 disease, clinical complications involving the metabolic/endocrine system are frequently observed. These include critical alterations of glycaemic control in diabetic patients and new-onset hyperglycaemia at admission in individuals without previous clinical history of diabetes. Although multiple causes have been indicated for COVID-19-related hyperglycaemia, a recently published case report described autoantibody-negative insulin-dependent diabetes onset in a young patient who was infected by SARS-CoV-2 seven weeks before occurrence of diabetes symptoms. This suggests a potential effect (direct or indirect) of the SARS-CoV-2 infection on the pancreatic islet insulin producing β-cells, but additional evidence is needed to allow solid conclusions.

Previous studies suggested that ACE2, the human host cell receptor for SARS-CoV-2 and SARS-CoV, which in other tissues has been shown to be a necessary component for infection permissiveness (6,50) is expressed in pancreatic tissue (19). However, an in-depth analysis aimed at evaluating ACE2 expression pattern distribution in human pancreas is still lacking. Here, we adopted multiple technologies and reagents to thoroughly analyse presence of ACE2, both at mRNA and protein level, in order to evaluate its expression and localization in pancreatic tissue samples obtained from adult non-diabetic multiorgan donors from the INNODIA EUnPOD biobank collection, in enzymatic- and LCM-isolated primary adult human pancreatic islets and in human β-cell line EndoC-βH1.

In human adult pancreas, we primarily observed ACE2 expression in microvasculature component (endothelial cells-associated pericytes, both in endocrine and exocrine compartments). The expression of ACE2 in the pancreatic microvasculature compartment was associated, but not superimposed, to the endothelial cells specific marker CD31 (or PECAM-1). Such staining pattern strongly suggests the presence of ACE2 in pancreatic vascular pericytes which are tightly associated to endothelial cells.

Of interest, although the exocrine pancreas and the pancreatic islets are highly vascularized (51), only a subset of pancreatic pericytes cells markedly express ACE2. Additionally, ACE2 expression in microvascular compartment is surprisingly lobular, resembling the heterogeneous staining pattern of several inflammatory markers. Additional observations on multiple pancreatic lobules are required to confirm such heterogeneous lobular patterning observed.

The presence of ACE2 in pancreatic pericytes is of sure interest. As a matter of fact, the vascular leakage and endothelitis were reported as a typical sign of SARS-CoV-2 infection in various organs, driving early local inflammation and the subsequent exacerbation of immune responses (52,53). Of note, multiple studies addressed the importance of an intact islet microvasculature in order to render pancreatic islets fully functional (54,55). Therefore, a pancreatic islet local vascular damage and inflammation due to SARS-CoV-2 direct infection of ACE2^+^ pancreatic pericyte cells is a potential contributory factor for islet dysfunction. Of note, two recent preprint manuscripts also indicated the presence of ACE2 expression in pancreatic microvasculature (56,57).

Our results indicate that ACE2 is expressed also in the pancreatic islets and this expression is mostly located in β-cells as compared to α-cells. These results are in contrast with the two recent preprint manuscripts (56,57) which failed to observe ACE2 expression in pancreatic islet endocrine cells. Such discrepancies may be explained by a resulting sum of differences in primary antibodies sensitivity, different epitopes targeted, tissue sections preparation and pre-treatment, as well as immunodetection methodology sensitivity (immunohistochemistry vs. immunofluorescence). Such variables may be of critical importance when detecting a low-expressed protein and may generate different results. Furthermore, it should be taken into consideration that ACE2 expression may vary greatly among individuals due to genetic or environmental factors (58). Such intrinsic ACE2 variability has been previously observed also in other cellular contexts, with some authors reporting high ACE2 levels and others low or absent (59).

In our study, localization of ACE2 in pancreatic islet endocrine cells was observed using two out of three different antibodies tested. Surprisingly, we were not able to observe ACE2 pancreatic islets positive signal using Abcam monoclonal antibody Ab108252, while signal was clearly evident in microvasculature and scattered ductal cells. However, these results are in line with Kusmartseva et al 2020 (57). Of note, Ab108252 monoclonal antibody specifically targets an epitope located in the N-terminal domain of ACE2 protein (200-300aa) which is missing in the recently discovered truncated ACE2 isoform (short-ACE2, 357-805aa) (24,25), thus being capable to recognize only the long-ACE2 isoform. On the contrary, by using two different antibodies (MAB933 and Ab15348) - which can recognize the C-terminal domain shared between short- and long-ACE2 - we obtained clear and concordant results, with identification of ACE2 in pancreatic islet endocrine cells in addition to the microvasculature. As a positive control for the antibodies and our IHC procedure, MAB933 and Ab15348 were also tested in FFPE lung tissue sections, showing overlapping results with previously published studies (7,41,42).

Overall, these results suggest that the short-ACE2 isoform may be the prevalent one expressed in β-cells. Indeed, ACE2 western blot analyses of the human β-cell line EndoC-βH1 support this hypothesis by confirming that: (*i*) short-ACE2 is prevalent over long-ACE2, the latter being present but with low expressed in β-cells; (*ii*) Abcam Ab108252 cannot recognize the short-ACE2 isoform. Based on these results we suggest that in human β-cells both ACE2 isoforms are present, with a predominance of the short-ACE2.

Although the presence of the short-ACE2 isoform, alongside with long-ACE2, is clearly evident in human β-cells, its functional role remains to be established. A previous study suggested the ability of the short-ACE2 isoform to form homodimers or heterodimer with the long-ACE2 isoform, thus potentially being able to modulate both the activity and structural protein domains conformation of the long-ACE2 (22). Significantly, short-ACE2 is missing the aminoacidic residues reported as fundamental for virus binding; however, due to the lack of detailed data regarding its function we cannot exclude that the short-ACE2 may modulate SARS-CoV-2 susceptibility by interacting with long-ACE2 or additional membrane proteins which may mediate the binding to SARS-CoV-2 spike protein [e.g. ITGA5, as previously observed (60)]. Of interest, nasal epithelium, reported to represent the main reservoir of SARS-CoV-2 (61), showed higher levels of short-ACE2 vs. long-ACE2 (24).

As described above, we report that in human pancreatic islets ACE2 is enriched in insulin-producing β-cells. This *in-situ* ACE2 expression pattern data are in line with three different bulk RNA-seq datasets analysing human β- and α-cells transcriptome, reporting higher expression of ACE2 mRNA in β-cells as compared to α-cells (62–64) (**Figure S13D-F**). Of note, such datasets rely on β- or α-cells bulk RNA-seq and do not suffer from common limitations present in single-cell RNA seq (65), which usually detects only 1,000-3,000 genes/cell, thus representing a minor fraction (25-30%) of the total number of genes identified by bulk cells RNA sequencing (>20,000 genes). Another recent study analyzed SARS-CoV-2 host receptors expression using two different methods (microarray and bulk RNA-seq) and further confirmed that ACE2 is indeed expressed in human pancreatic islets, and also demonstrated that ACE2 expression was higher in sorted pancreatic β-cells relative to other endocrine cells (66). Additional evidence of ACE2 expression in endocrine pancreas and in β-cells derive also from mouse studies, which demonstrated ACE2 expression in insulin-producing cells as well as its critical role in the regulation of β-cell phenotype and function (67–70).

Collectively, our in-situ expression data alongside with multiple published datasets and reports both in human and mouse show that ACE2 is expressed in pancreatic islets, albeit at relatively low levels. ACE2 expression in human β-cells may render these cells sensitive to SARS-CoV-2 entry (23). Such hypothesis is consistent with the known sensitivity of these cells to infection by several enterovirus serotypes. Indeed, multiple evidence from our and other groups (71–74) showed that enteroviruses are capable to competently infect β-cells but less so α-cells (75,76); these viruses are thus being considered as one of the potential triggering causes of type 1 diabetes (T1D) (77). Of note, it has been previously demonstrated that human β-cells exclusively express virus receptor isoform Coxsackie and adenovirus receptor-SIV (CAR-SIV), making them prone to infection by certain viruses (78). Therefore, it would not be surprising that, under particular conditions, human β-cells could be directly infected by SARS-CoV-2. Importantly, a recent report showed that human pancreatic islets can be infected *in vitro* by SARS-CoV-2 (23), supporting our observations of a specific tropism of the virus due to ACE2 expression.

Noteworthy, the subcellular localization of ACE2 in β-cells recapitulates what was previously found for the virus receptor CAR-SIV (78). In our dataset, ACE2 protein signal is mostly cytoplasmic/granular and partially overlaps with insulin granules. Additional spots are also localized close to the plasma membrane, thus suggesting the existence of ACE2 isoforms in multiple compartments within β-cells. Such subcellular localization was observed both *in-vitro* in EndoC-βH1 and *ex vivo* in β-cells of primary human pancreatic tissues. Although ACE2 has been primarily observed on cell surfaces (42,79), some studies also described ACE2 granular localization in other cell types of epithelial origin (9,80). A similar intracellular localization and putative trafficking was observed for the viral receptor CAR-SIV (78), also expected to be mainly localized on the cell membrane. Against this background, we speculate that: (i) upon activation, ACE2 can be internalized through endosome/lysosome pathway (81); (ii) in β-cells, ACE2 trafficking to cell membrane may be mediated by insulin granules; (iii) in β-cells the short-ACE2 isoform could be mainly localized in cytoplasm, while long-ACE2 in the plasma membrane; (iii) ACE2 can be secreted and found in a soluble form or within exosomes (82,83). Additional studies are needed to fully ascertain the subcellular localization and trafficking of ACE2 isoforms in human β-cells, and to determine whether ACE2 is indeed present in the secretome of β-cells.

Our data also indicate that in β-cells, total ACE2 mRNA expression is upregulated upon different pro-inflammatory conditions, but not following exposure to the metabolic stressor palmitate or to the T2D environment. Importantly, these observations were obtained both in the human β-cell line EndoC-βH1 and in human primary pancreatic islet cells, as shown by qRT-PCR, RNA-seq datasets and immunofluorescence. As a matter of fact, ACE2 has been previously indicated as an interferon-stimulated gene (ISG) in a variety of cells (58). Of interest, a previous report suggested that the short-ACE2 may represent the prevalent ACE2 isoform upregulated upon inflammatory stress (25). However, whether this is the case also in human β-cells it should be examined in depth in future studies. Although total ACE2 expression is increased upon inflammatory stress, in-situ analysis of ACE2 in infiltrated pancreatic islets derived from FFPE sections of a pancreas from a longstanding T1D donor did not reveal significant changes of SARS-CoV2 receptor expression. However, we recognize that a high number of pancreatic islets with different degree of inflammation, and pancreata from recent-onset T1D donors showing highly infiltrated islets are needed to adequately characterize ACE2 expression in the early stages of the disease and to determine whether changes in ACE2 expression contribute to: (*i*) the observed alteration of glycaemic control at admission in SARS-CoV-2 individuals without previous clinical history of diabetes; (*ii*) the increased severity of COVID-19 in those subjects with previous inflammatory-based diseases.

In conclusion, the presently described preferential expression of ACE2 isoforms in human β-cells, alongside with ACE2 upregulation under pro-inflammatory conditions *in-vitro*, identifies the putative molecular basis of SARS-CoV-2 tropism for pancreatic islets and uncovers a link between inflammation and ACE2 expression levels in islet β-cells. Further mechanistic studies and epidemiological evaluation of individuals affected by COVID-19, including long-term clinical follow up and determination of islet autoantibodies, may help to clarify whether human β-cells are among the target cells of SARS-CoV-2 virus and whether β-cell damage, with the potential induction of autoimmunity in genetically susceptible individuals, occurs during and/or after infection.

## Supporting information

Supplementary Material_ Fignani et al 2020

## 6 Author Contributions

GS, FD, DF, GL, NB conceived and designed the experiments. GS, DF, GL and NB performed RT-Real Time PCR, Laser Capture Microdissection and Immunohistochemical/Confocal fluorescence Microscopy analysis experiments. DF and GEG performed EndoC-βH1 cell culture experiments. PM and LM isolated human pancreatic islets and contributed to the collection and processing of human pancreata. LN and NB curated FFPE processing and data analysis of INNODIA EUnPOD pancreata biobank collection. DLE and MC conceived and performed RNA-seq on EndoC-βH1 and human pancreatic islets and contributed to the scientific discussion. CM, CG and LO conceived the experiments and contributed to the scientific discussion. All authors reviewed and provided input to the manuscript.

## 8 Acknowledgments

We thank Dr. R. Scharfmann (Institut Cochin, INSERM, Université de Paris) who provided the human pancreatic β-cell line EndoC-βH1. We thank Dr. Laura Salvini, Dr. Laura Tinti and Dr. Vittoria Cicaloni (Toscana Life Sciences, Siena-Italy) for their expertise and support in the analysis of the shotgun proteomics data in EndoC-βH1. The secretarial help of Maddalena Prencipe and Alessandra Mechini (University of Siena) is greatly appreciated. This manuscript has been released as a pre-print at https://www.biorxiv.org/content/10.1101/2020.07.23.208041v1 (84).

