## Supplementary Material_ Fignani et al 2020 for "SARS-CoV-2 receptor Angiotensin I-Converting Enzyme type 2 (ACE2) is expressed in human pancreatic β-cells and in the human pancreas microvasculature"

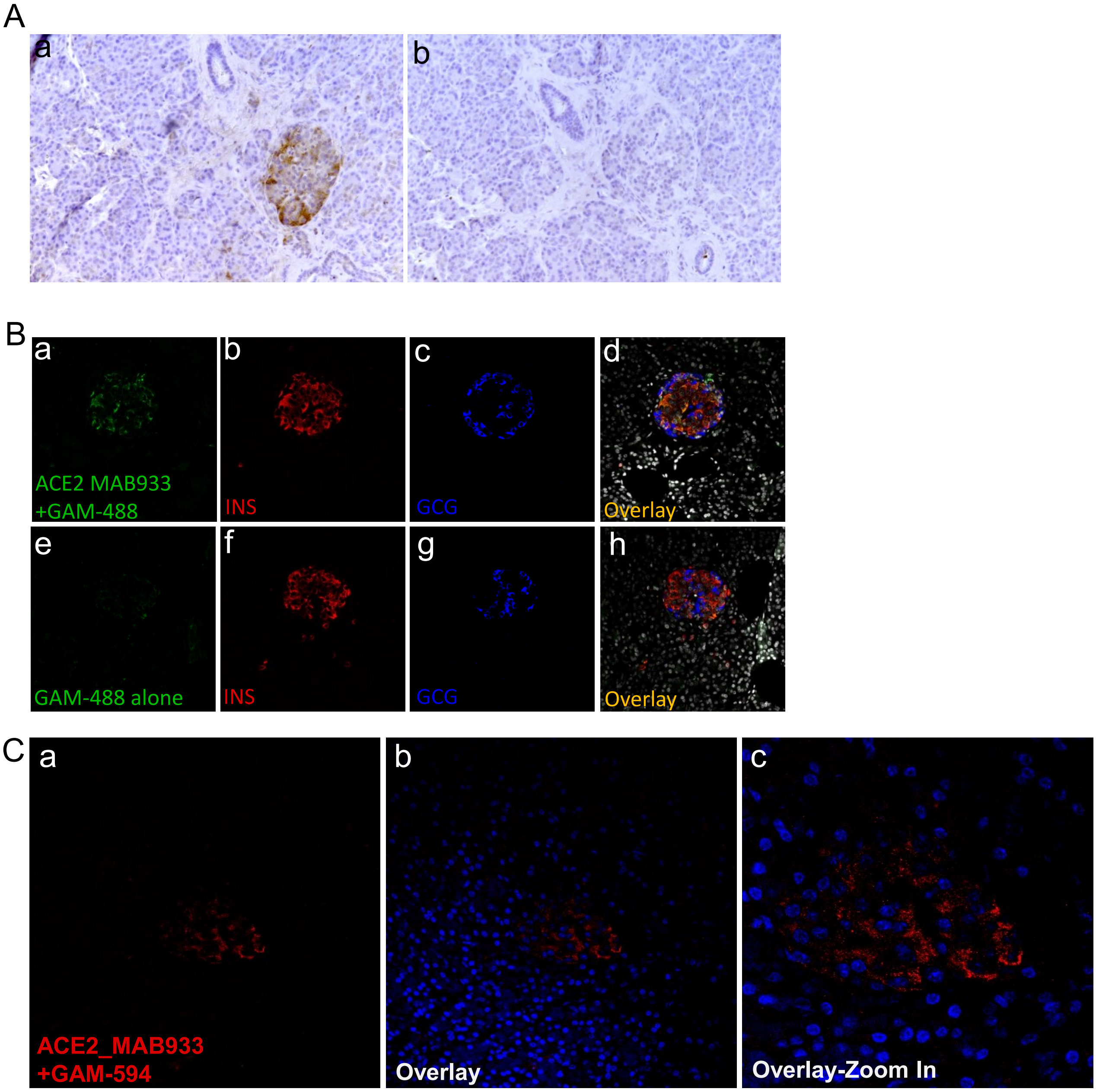

**Figure S1. ACE2 negative control staining in human pancreas.** (**A**) Immunohistochemistry of a Formalin-Fixed Paraffin Embedded (FFPE) pancreatic section from an adult non-diabetic multiorgan donor (id: #210518) using MAB933 anti-ACE2 antibody (panel-a) or secondary antibody-only (panel-b). (**B**) Triple Immunofluorescence of a FFPE pancreatic section from an adult-non diabetic multiorgan donor (id: 210518) using MAB933 anti-ACE2 antibody (panel-a) or fluorescently-labeled Goat Anti Mouse (GAM) secondary antibody-only as negative control for ACE2 (panel-b). (**C**) ACE2 immunofluorescent staining in FFPE pancreatic section from an adult-non diabetic multiorgan donor (id: 210518) using MAB933 anti-ACE2 antibody (panel-a) with another fluorescently labeled Goat Anti Mouse-594, in order to exclude background staining artifacts due to the previously adopted ACE2 fluorescent detection system.

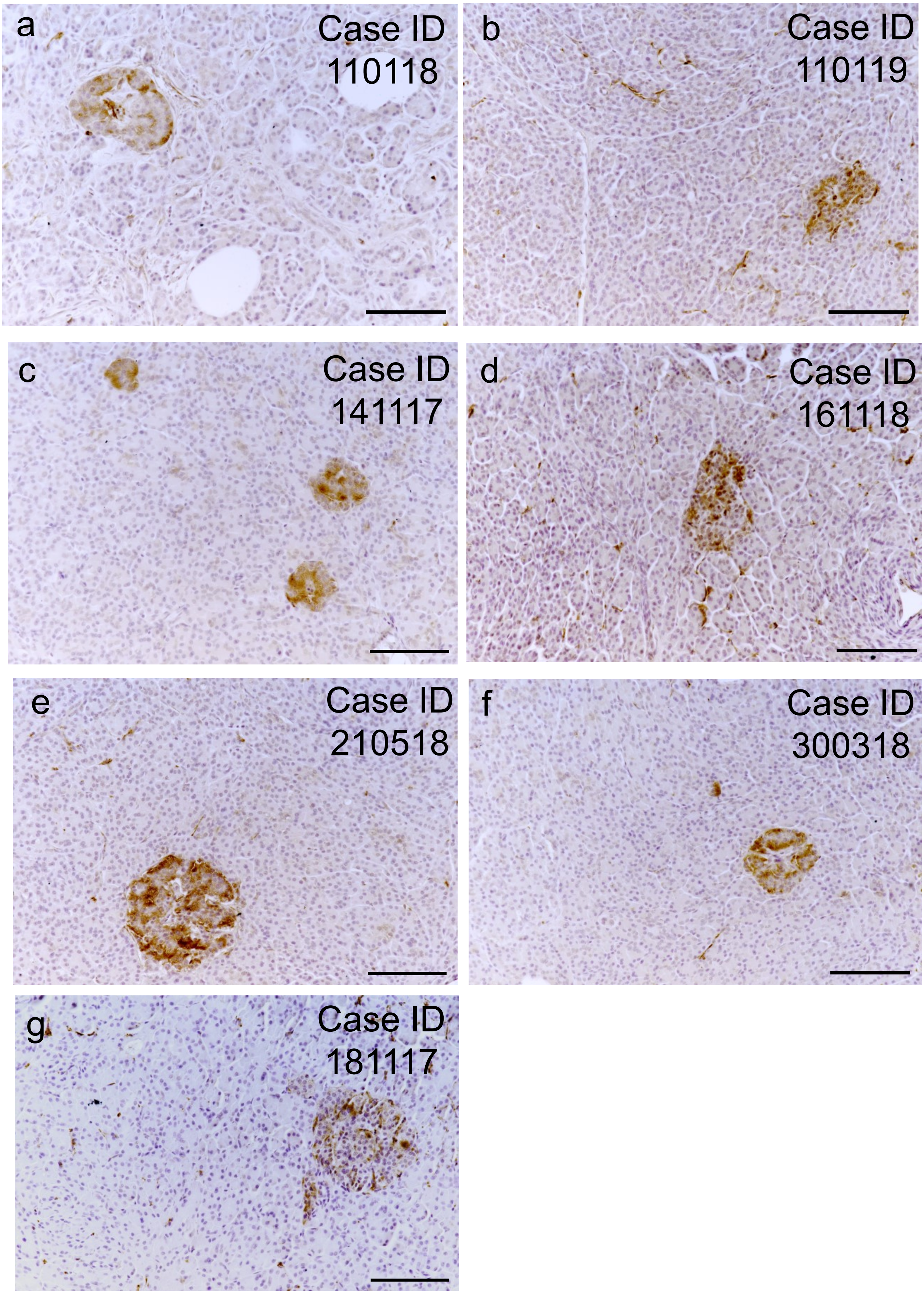

**Figure S2.** Representative images of all cases showing expression of ACE2 (MAB933) in human pancreatic islets. Representative images of ACE2 IHC staining in human pancreatic FFPE sections obtained from seven different non-diabetic multiorgan donors. Scale bar: 100 µm.

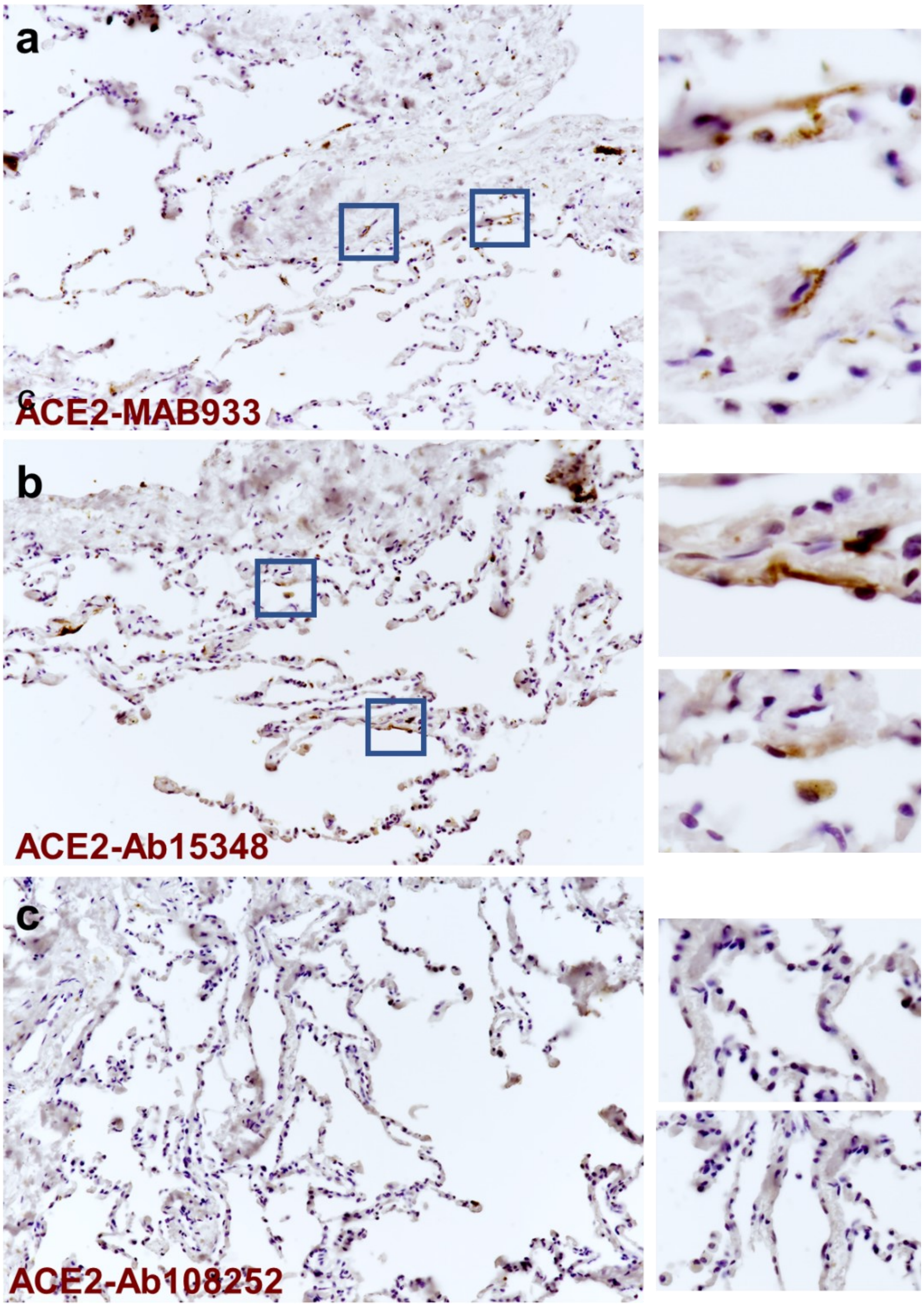

**Figure S3**. **ACE2 staining positive control**. Anti-ACE2 MAB933 (panel-a), Ab15348 (panel-b) and Ab108252 (panel-c) antibodies immunohistochemistry in FFPE lung tissue sections. Zoom-in insets are shown on the right, adjacent to each main related image.

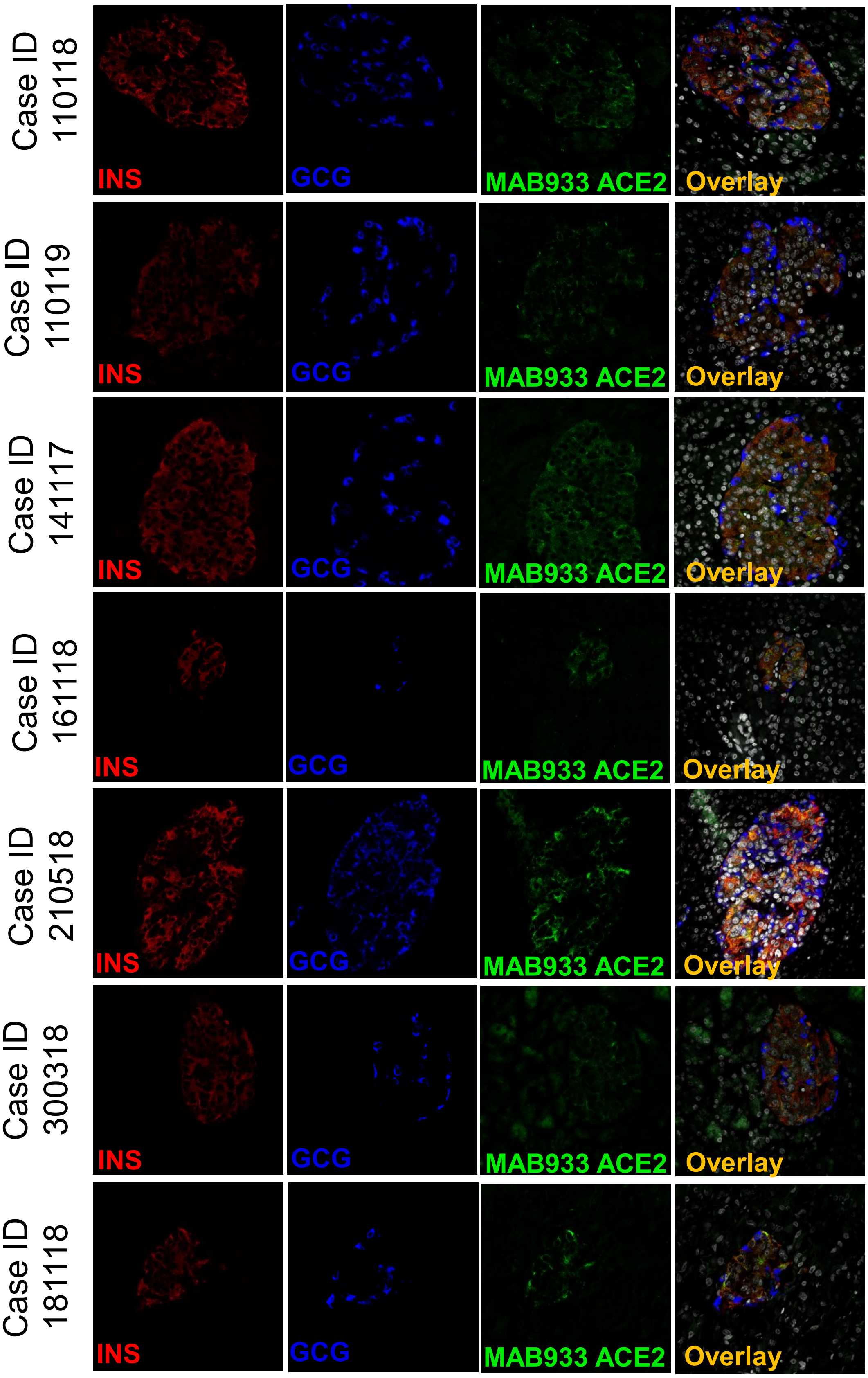

**Figure S4. ACE2 MAB933 immunofluorescence in FFPE pancreatic sections**. Representative images of all cases showing expression of insulin (red), glucagon (blue) and ACE2 (MAB933, green), in human pancreatic FFPE sections obtained from seven different adult non-diabetic multiorgan donors.

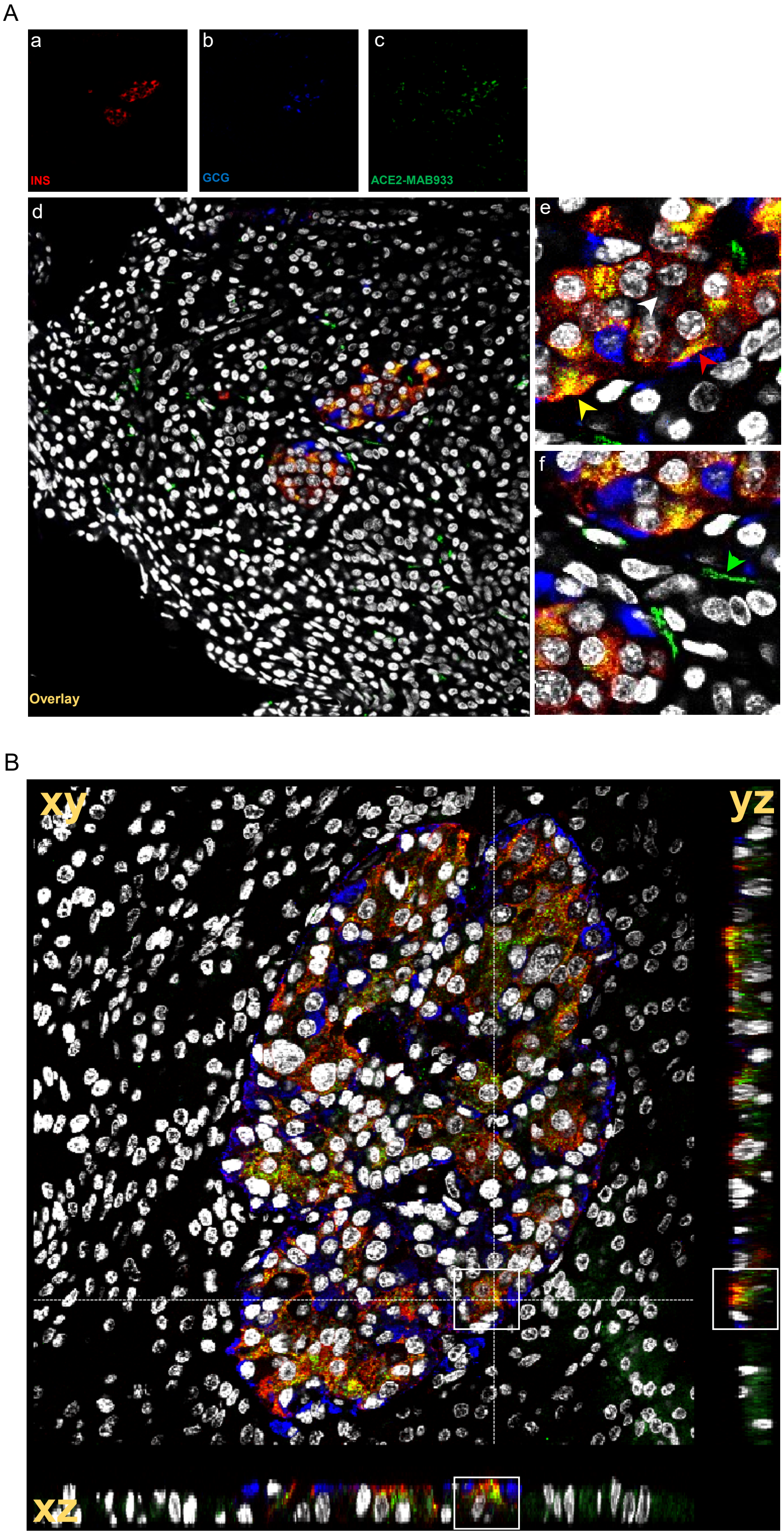

**Figure S5.** (**A**) Triple immunofluorescence staining and image analysis of FFPE human pancreatic section (case #210518) stained for insulin (red, panel-a), glucagon (blue, panel-b) and ACE2 MAB933 (green, panel-c), alongside with overlay (panel-d) and zoom overlays (panel-e and -f). White-arrow: ACE2-negative β-cell; yellow-arrow: granular/cytoplasmatic ACE2 signal in a β-cell;

red-arrow: ACE2-positivity prevalently associated to plasma membrane in a β-cell; green-arrow: plasma membrane localization of ACE2 in microvasculature-associated pericyte.

**(B) Z-stack 3D analysis of a pancreatic islet stained for ACE2 (MAB933), insulin and glucagon.** Representative image of a human pancreatic islet of case 210518 analysed performing triple immunofluorescence staining for ACE2 (green), insulin (red), glucagon (blue) and DAPI (white). Acquisition of 40 different focal planes allowed a XYZ 3D reconstruction and z-sectioning (xz and yz). White rectangles highlight an example of the partial colocalization of ACE2 and insulin signals within β-cells, highlighting the different subcellular compartmentalization (cytoplasmatic/granular and plasma membrane) of ACE2.

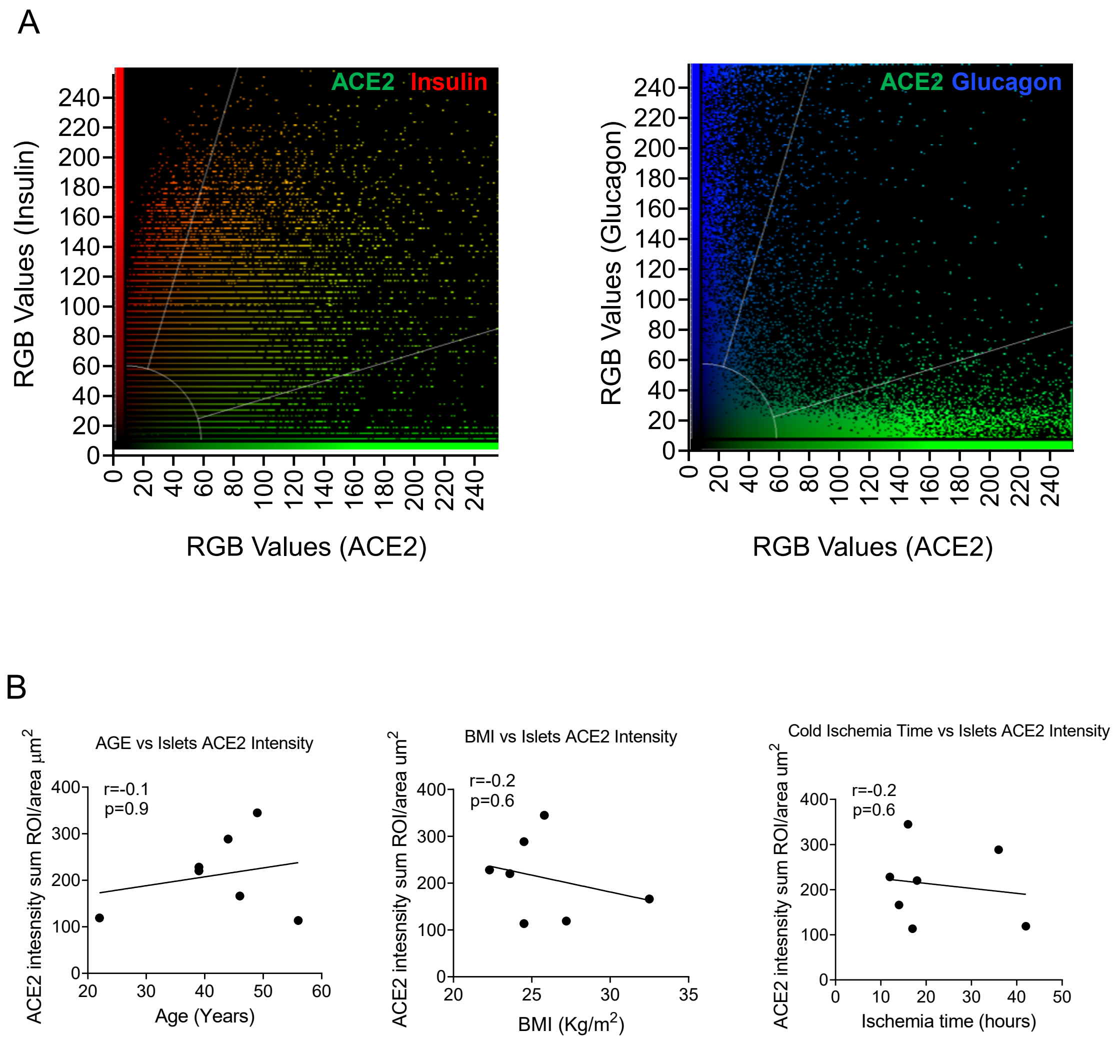

**Figure S6. (A)** Colocalization plots of ACE2-insulin and ACE2-glucagon in a human pancreatic islet of case 210518. (**B**) Correlation analysis among ACE2 pancreatic islets staining intensity of all cases analyzed (see Figure 4C) and Age (years), BMI (kg/$m^2$) and cold ischemia time (hours). Statistics performed using Spearman R correlation test.

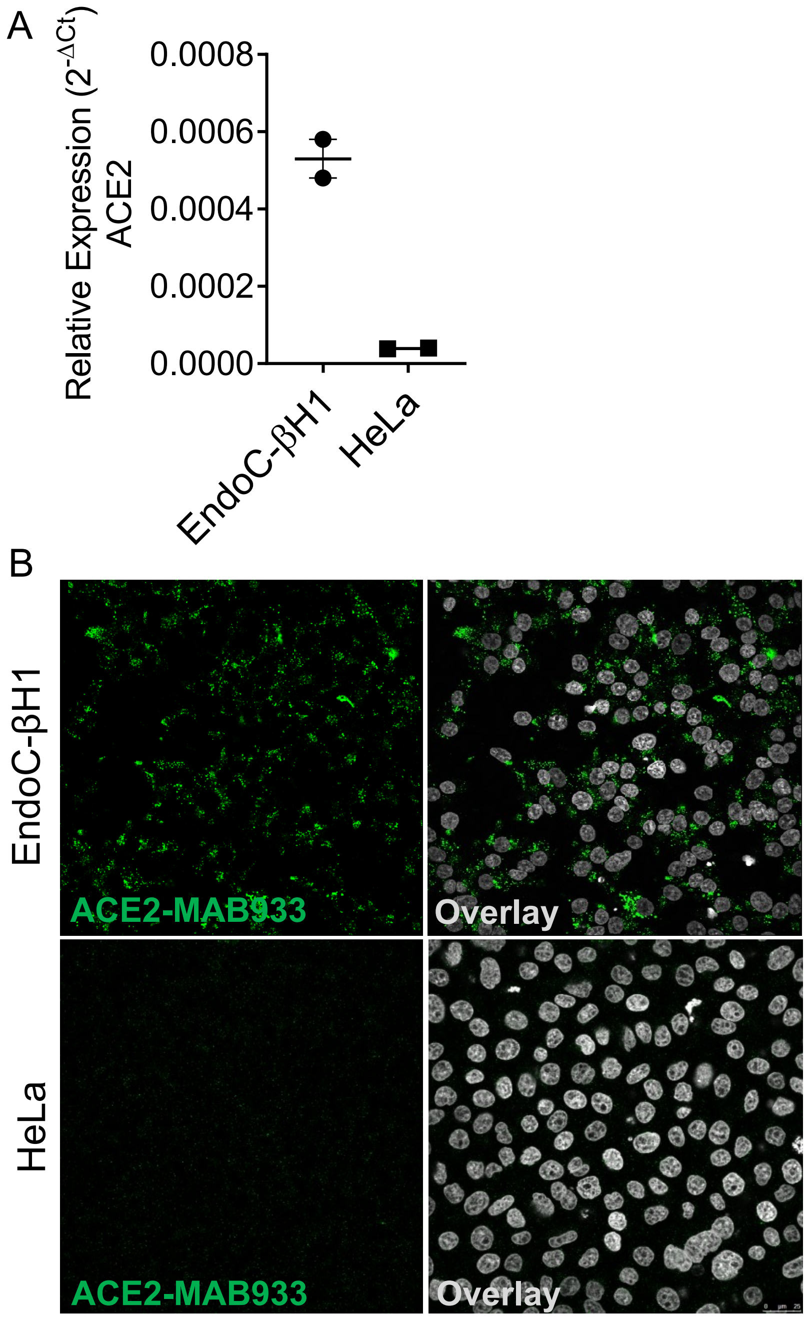

**Figure S7. (A) Orthogonal validation of ACE2 R&D MAB933 antibody in cultured cell lines.** ACE2 mRNA expression using qRT-PCR in EndoC-βH1 and HeLa cells. Cycle Threshold values of ACE2 were normalized using β-Actin and GAPDH values. Final values are reported as normalized $2^{-dCt}$ relative values. (B) Immunofluorescence analysis of ACE2 (R&D, MAB933) in EndoC-βH1 and HeLa cells stained using the same protocol.

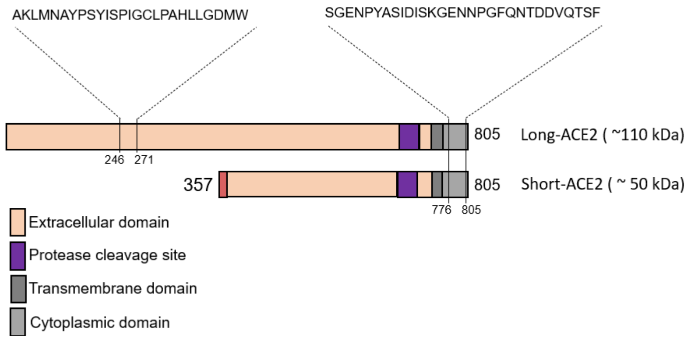

```
# Aligned_sequences: 2
# 1: EMBOSS_001
# 2: EMBOSS_001
# Matrix: EBLOSUM62
# Gap_penalty: 10.0
# Extend_penalty: 0.5
#
# Length: 805
# Identity:     452/805 (56.1%)
# Similarity:   452/805 (56.1%)
# Gaps:         346/805 (43.0%)
# Score: 2387.0
#

EMBOSS_001         1 MSSSSWLLLSLVAVTAAQSTIEEQAKTFLDKFNHEAEDLFYQSSLASWNY     50

EMBOSS_001         1 --------------------------------------------------      0

EMBOSS_001        51 NTNITEENVQNMNNAGDKWSAFLKEQSTLAQMYPLQEIQNLTVKLQLQAL    100

EMBOSS_001         1 --------------------------------------------------      0

EMBOSS_001       101 QQNGSSVLSEDKSKRLNTILNTMSTIYSTGKVCNPDNPQECLLLEPGLNE    150

EMBOSS_001         1 --------------------------------------------------      0

EMBOSS_001       151 IMANSLDYNERLWAWESWRSEVGKQLRPLYEEYVVLKNEMARANHYEDYG    200

EMBOSS_001         1 --------------------------------------------------      0

EMBOSS_001       201 DYWRGDYEVNGVDGYDYSRGQLIEDVEHTFEEIKPLYEHLHAYVRAKLMN    250

EMBOSS_001         1 --------------------------------------------------      0

EMBOSS_001       251 AYPSYISPIGCLPAHLLGDMWGRFWTNLYSLTVPFGQKPNIDVTDAMVDQ    300

EMBOSS_001         1 --------------------------------------------------      0

EMBOSS_001       301 AWDAQRIFKEAEKFFVSVGLPNMTQGFWENSMLTDPGNVQKAVCHPTAWD    350
                                                                .....||
EMBOSS_001         1 -------------------------------------------MREAGWD      7

EMBOSS_001       351 LGKGDFRILMCTKVTMDDFLTAHHEMGHIQYDMAYAAQPFLLRNGANEGF    400
                     .|.   ||||||||||||||||||||||||||||||||||||||||||||
EMBOSS_001         8 KGG---RILMCTKVTMDDFLTAHHEMGHIQYDMAYAAQPFLLRNGANEGF     54

EMBOSS_001       401 HEAVGEIMSLSAATPKHLKSIGLLSPDFQEDNETEINFLLKQALTIVGTL    450
                     ||||||||||||||||||||||||||||||||||||||||||||||||||
EMBOSS_001        55 HEAVGEIMSLSAATPKHLKSIGLLSPDFQEDNETEINFLLKQALTIVGTL    104

EMBOSS_001       451 PFTYMLEKWRWMVFKGEIPKDQWMKKWWEMKREIVGVVEPVPHDETYCDP    500
                     ||||||||||||||||||||||||||||||||||||||||||||||||||
EMBOSS_001       105 PFTYMLEKWRWMVFKGEIPKDQWMKKWWEMKREIVGVVEPVPHDETYCDP    154

EMBOSS_001       501 ASLFHVSNDYSFIRYYTRTLYQFQFQEALCQAAKHEGPLHKCDISNSTEA    550
                     ||||||||||||||||||||||||||||||||||||||||||||||||||
EMBOSS_001       155 ASLFHVSNDYSFIRYYTRTLYQFQFQEALCQAAKHEGPLHKCDISNSTEA    204

EMBOSS_001       551 GQKLFNMLRLGKSEPWTLALENVVGAKNMNVRPLLNYFEPLFTWLKDQNK    600
                     ||||||||||||||||||||||||||||||||||||||||||||||||||
EMBOSS_001       205 GQKLFNMLRLGKSEPWTLALENVVGAKNMNVRPLLNYFEPLFTWLKDQNK    254

EMBOSS_001       601 NSFVGWSTDWSPYADQSIKVRISLKSALGDKAYEWNDNEMYLFRSSVAYA    650
                     ||||||||||||||||||||||||||||||||||||||||||||||||||
EMBOSS_001       255 NSFVGWSTDWSPYADQSIKVRISLKSALGDKAYEWNDNEMYLFRSSVAYA    304

EMBOSS_001       651 MRQYFLKVKNQMILFGEEDVRVANLKPRISFNFFVTAPKNVSDIIPRTEV    700
                     ||||||||||||||||||||||||||||||||||||||||||||||||||
EMBOSS_001       305 MRQYFLKVKNQMILFGEEDVRVANLKPRISFNFFVTAPKNVSDIIPRTEV    354

EMBOSS_001       701 EKAIRMSRSRINDAFRLNDNSLEFLGIQPTLGPPNQPPVSIWLIVFGVVM    750
                     ||||||||||||||||||||||||||||||||||||||||||||||||||
EMBOSS_001       355 EKAIRMSRSRINDAFRLNDNSLEFLGIQPTLGPPNQPPVSIWLIVFGVVM    404

EMBOSS_001       751 GVIVVGIVILIFTGIRDRKKKNKARSGENPYASIDISKGENNPGFQNTDD    800
                     ||||||||||||||||||||||||||||||||||||||||||||||||||
EMBOSS_001       405 GVIVVGIVILIFTGIRDRKKKNKARSGENPYASIDISKGENNPGFQNTDD    454

EMBOSS_001       801 VQTSF    805
                     |||||
EMBOSS_001       455 VQTSF    459
```

**Figure S8. ACE2 Targeted Mass Spectrometry-Shotgun Proteomic Analysis of EndoC-βH1.** **(A)** Representative scheme of ACE2 isoforms (short-ACE2 and long-ACE2) and consistent peptides identified in EndoC-βH1 using shotgun proteomics in n=2 independent experiments. Below**,** alignment of ACE2 isoforms protein FASTA sequences performed using EMBOSS. Red sequence: short-ACE2; black sequence: long-ACE2; yellow highlighted sequences: shotgun proteomics identified peptides**.**

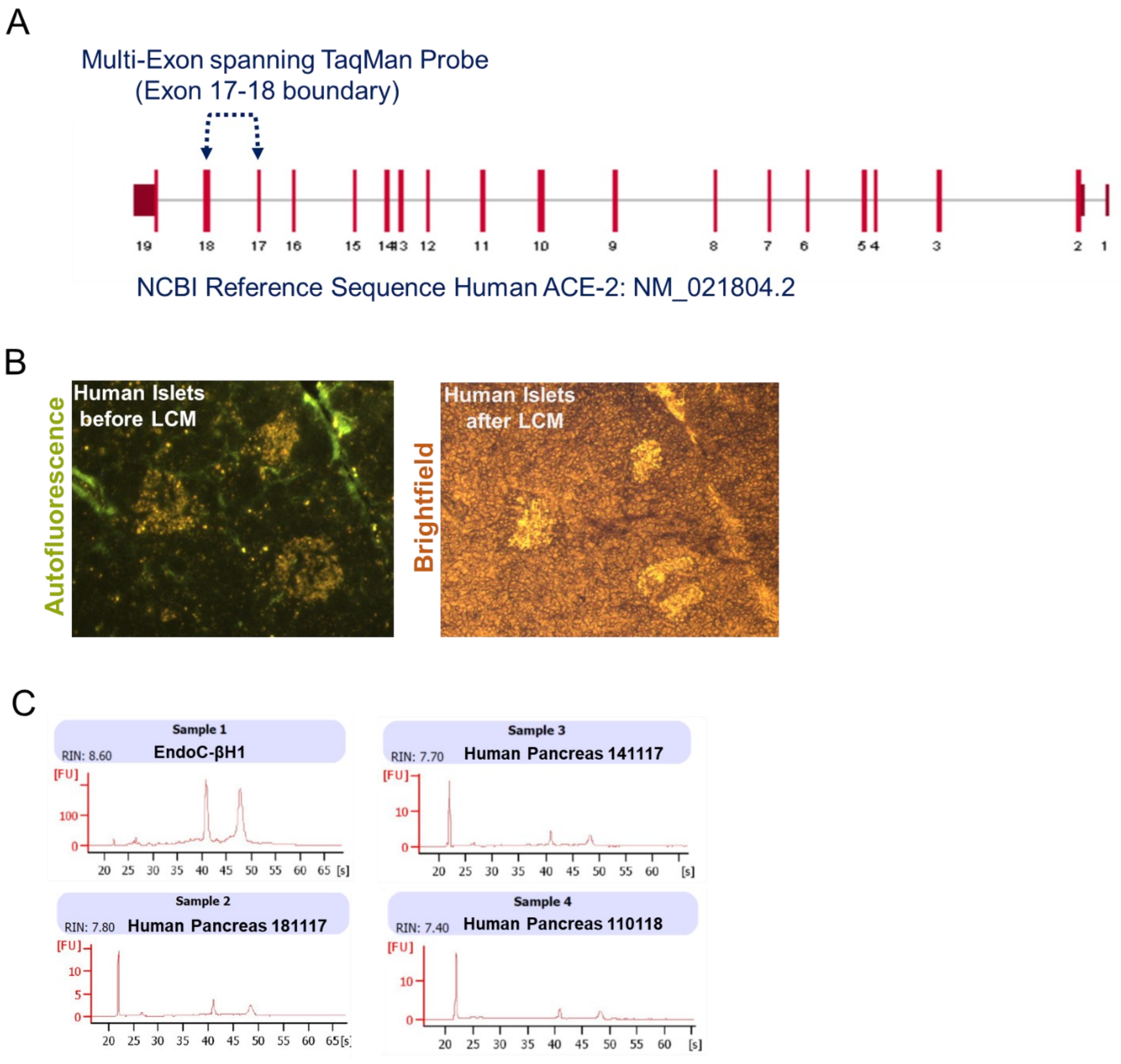

**Figure S9. ACE2 mRNA expression. (A)** ACE2 gene scheme reporting exons of full length transcript (Ref Seq: NM_021804.2) and showing the location of the TaqMan Assay (Hs01085333_m1, assay location 2332nt, exon 17-18, amplicon length:141nt) used to detect ACE2 mRNA expression. (**B**) Human pancreatic islets autofluorescence and brightfield images in a human pancreatic frozen section, before and after laser capture microdissection procedure. (C) Electropherogram of total RNA extracted from EndoC-βH1 cells and from LCM-islets from human pancreatic sections obtained from five non-diabetic multiorgan donors. Representative RNA Integrity Number (RIN) is reported for each RNA sample analyzed.

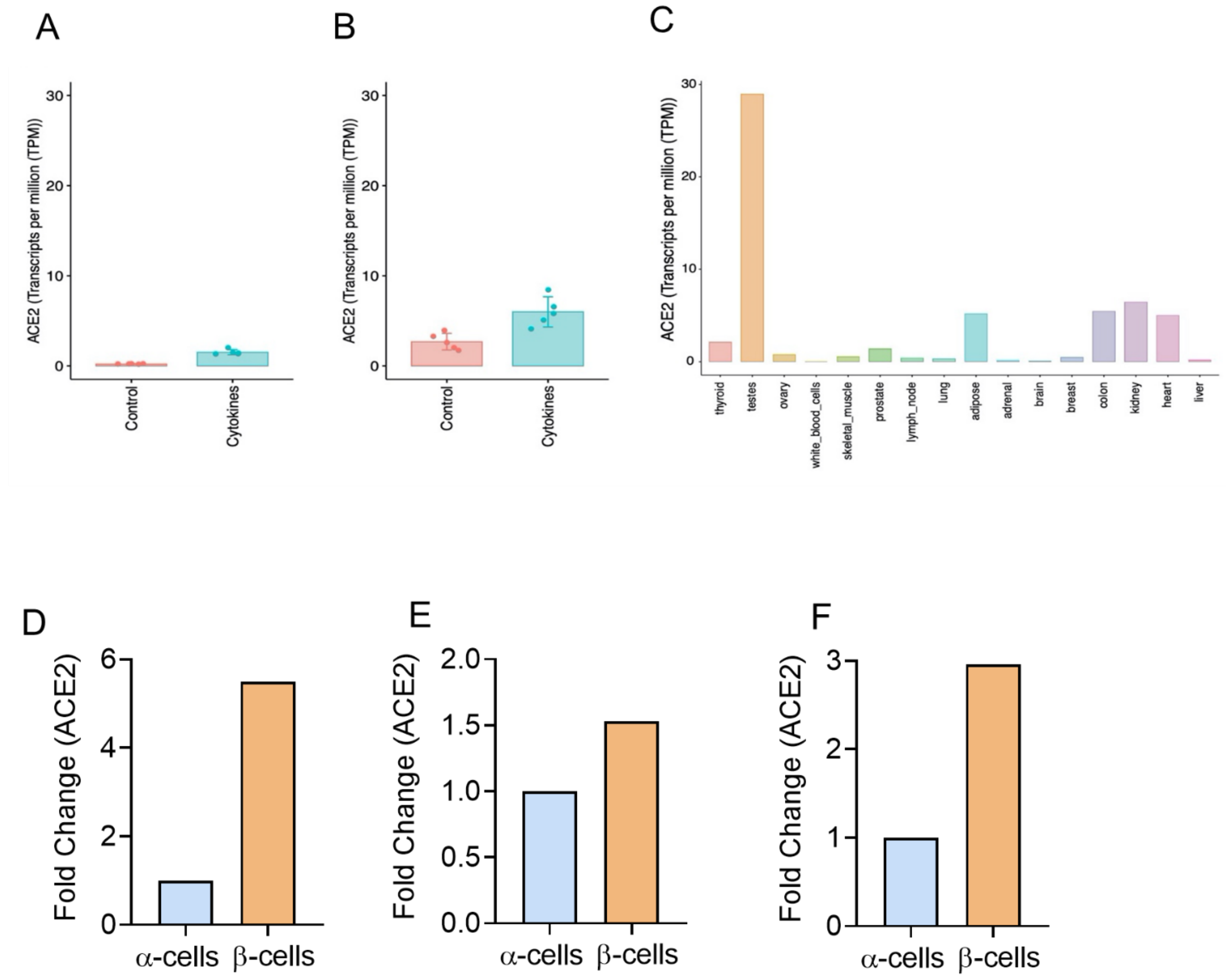

**Figure S10.** Expression of ACE2 mRNA expression in different conditions and tissues. The expression of ACE2 was evaluated in EndoC-βH1 cells (**A**) and pancreatic human islets (**B**) exposed or not to proinflammatory cytokines (IL1β + IFNγ) for 48h and in 16 different tissues from the Illumina Body Map 2.0 (GEO: GSE30554)) (**C**) that were sequenced at a similar deep coverage (coverage > 150 x 106 reads / per sample). Values are represented as transcripts per million (TPM). All the samples were processed using the same RNA-seq pipeline as described in Methods.

(D-E) ACE2 expression in β-cells and in α-cells, obtained from three different RNA-seq datasets. Data were extrapolated from supplementary files of normalized and analyzed datasets obtained from three different published studies which compared β- and α-cells transcriptomes. In (**D**) Dorrell et al 2011, in (**E**) Bramswig et al 2013, and in (**F**) Blodgett et al 2015 (Blodgett et al., 2015; Bramswig et al., 2013; Dorrell et al., 2011).

A

ACTGCTGTCCCAGGCTCCTTGTTCATTTATTCTTAGCTTTCAGATTTTATATCTTGCTTT
CTCCATTTCAGTTTGATCAAGTTAATCGTATTTTCTGCCTCAGCGAATTGAATTTGGACT
A**CTCCCATTTTTTCCACTAAACC**CACGTGTTTCATTTAGGAACTTACCTGTGTAGAAATT
STAT1

TATTTCCGCTTTTA**TTACAGTAA**CATTTCAACCTTTTCTTCAGATACTCGTGTGCTGTCC
STAT

TCTTTCTTGCCTGTTTTTTCTCCTATTGTGTGCAAGGAGTGAGT**GGATCTAGGTTTCCTG**
STAT1
STAT3
STAT5
STAT

**GAATGTGGGAGGA**GCTTTTCTGAAATTTCTGCACTGATTTTTTTTTTTTTACCCTCATCTC

ACTTTCATTTAAAATTCTTTATTATTATTACTTATATTTGTGAGCTGCTTTATTTTTTTA
AAAGGAGAGTATATTTTGATTCACATCAAGGGAAGTAAGAGTCCACCTCTTTTTGATGCA
GTGTAACTAAATCTGTCATCTTCACTGAGGCACAAGGCCCTCTGTCACATTTTATCTATG
CCTCCACTATTGCTTTTAGACTGTTGACTACAAGTTAGTTAAGAGCCTATACTCTAGAAT
CAGAATACATAGATCCATGTTCTGATTCCATCATTTGTTAGCTGAGTGAGATATAGCAAT
TTACTTAAACTTAGTTTCATAATTCATAAAATGGTGATAAAATATTTGCCTATAGGTTTC
TTGTGATGCTCAAATGAGATATTATAACACAAAGCACTTTGAATAGTTACTTATAATTGT
TTTTTTCCTATTCTACAATAATATACACAACTTTGGAATAAGGAAAAGCAGTGGACATTT
TTTTAAAGGCTTGATTATTGCAATGTCACCTGAACCTGGAAGACTTGTTTTTCTGGGTGA
AGAAATATTTTCTCTGTGTCAGAGTTTCACAATCATCGTCAGGTAGGCCCTTGAACCCTG
CCATTTAAAGTGCTCCTCTCTTTGATCTGTGGCACTCATA

| Rank | P-value | Matrix_ID | Matrix_name |
|---|---|---|---|
| 1 | 0,0009 | M01478 | V$CPHX_01 |
| 2 | 0,0043 | M00412 | V$AREB6_01 |
| 3 | 0,0063 | M00250 | V$GFI1_01 |
| 4 | 0,0084 | M01058 | V$GFI1B_01 |
| 5 | 0,0100 | M01232 | V$SATB1_01 |
| **6** | **0,0117** | **M00457** | **V$STAT5A_01** |
| **7** | **0,0118** | **M01260** | **V$STAT1_05** |
| 8 | 0,0119 | M00119 | V$MAX_01 |
| **9** | **0,0124** | **M00459** | **V$STAT5B_01** |
| 10 | 0,0129 | M00347 | V$GATA1_06 |
| 11 | 0,0129 | M01004 | V$HELIOSA_02 |
| 12 | 0,0157 | M00711 | V$ZTA_Q2 |
| 13 | 0,0160 | M00317 | V$LDSPOLYA_B |
| 14 | 0,0180 | M01446 | V$BARHL2_01 |
| 15 | 0,0190 | M01022 | V$LEF1_Q2_01 |
| 16 | 0,0191 | M01116 | V$CLOCKBMAL_Q6 |
| 17 | 0,0196 | M00615 | V$MYCMAX_03 |
| 18 | 0,0198 | M01425 | V$HNF1B_01 |
| 19 | 0,0201 | M01014 | V$SOX_Q6 |
| 20 | 0,0214 | M00632 | V$GATA4_Q3 |
| 21 | 0,0216 | M00318 | V$LPOLYA_B |
| 22 | 0,0217 | M00802 | V$PIT1_Q6 |
| 23 | 0,0231 | M01665 | V$IRF8_Q6 |
| **24** | **0,0251** | **M00223** | **V$STAT_01** |
| 25 | 0,0251 | M01352 | V$NKX29_01 |
| 26 | 0,0311 | M01124 | V$OCT4_02 |
| 27 | 0,0323 | M00302 | V$NFAT_Q6 |
| 28 | 0,0335 | M01185 | V$BCL6_02 |
| 29 | 0,0346 | M00033 | V$P300_01 |
| 30 | 0,0367 | M00799 | V$MYC_Q2 |
| 31 | 0,0376 | M00055 | V$NMYC_01 |
| 32 | 0,0384 | M01009 | V$HES1_Q2 |
| 33 | 0,0391 | M00451 | V$NKX3A_01 |
| 34 | 0,0399 | M01383 | V$NKX3A_02 |
| 35 | 0,0405 | M00671 | V$TCF4_Q5 |
| 36 | 0,0407 | M00320 | V$MTATA_B |
| 37 | 0,0408 | M00118 | V$MYCMAX_01 |
| 38 | 0,0423 | M01464 | V$HOXA10_01 |
| 39 | 0,0430 | M00123 | V$MYCMAX_02 |
| 40 | 0,0441 | M00472 | V$FOXO4_01 |
| 41 | 0,0458 | M01342 | V$CDP_03 |
| 42 | 0,0459 | M00059 | V$YY1_01 |
| **43** | **0,0460** | **M01595** | **V$STAT3_03** |
| 44 | 0,0469 | M01457 | V$NKX23_01 |

B

ACTGCTGTCCCAGGCTCCTTGTTCATTTATTCTTAGCTTTCAGATTTTATATCTTGCTTT
CTCCATTTCAGTTTGATCAAGTTAATCGTATTTTCTGCCTCAGCGAATTGAATTTGGACT
ACTCCCAT**TTTTTCCACTAAACC**CACGTGTTTCATTTAGGAACTTACCTGTGTAGAAATT
STAT1

TATTTCCGCTTTTATTACAGTAACATTTCAACCTTTTCTTCAGATACTCGTGTGCTGTCC

TCTTTCTTGCCTGTTTTTTCTCCTATTGTGTGCAAGGAGTGAGTGGATCTA**GGTTTCCTG**
STAT1
STAT3

**GAATGTGGGAGGA**GCTTTTCTGAAATTTCTGCACTGATTTTTTTTTTTTTACCCTCATCTC

ACTTTCATTTAAAATTCTTTATTATTATTACTTATATTTGTGAGCTGCTTTATTTTTTTA
AAAGGAGAGTATATTTTGATTCACATCAAGGGAAGTAAGAGTCCACCTCTTTTTGATGCA
GTGTAACTAAATCTGTCATCTTCACTGAGGCACAAGGCCCTCTGTCACATTTTATCTATG
CCTCCACTATTGCTTTTAGACTGTTGACTACAAGTTAGTTAAGAGCCTATACTCTAGAAT
CAGAATACATAGATCCATGTTCTGATTCCATCATTTGTTAGCTGAGTGAGATATAGCAAT
TTACTTAAACTTAGTTTCATAATTCATAAAATGGTGATAAAATATTTGCCTATAGGTTTC
TTGTGATGCTCAAATGAGATATTATAACACAAAGCACTTTGAATAGTTACTTATAATTGT
TTTTTTCCTATTCTACAATAATATACACAACTTTGGAATAAGGAAAAGCAGTGGACATTT
TTTTAAAGGCTTGATTATTGCAATGTCACCTGAACCTGGAAGACTTGTTTTTCTGGGTGA
AGAAATATTTTCTCTGTGTCAGAGTTTCACAATCATCGTCAGGTAGGCCCTTGAACCCTG
CCATTTAAAGTGCTCCTCTCTTTGATCTGTGGCACTCATA

| Rank | P-value | Matrix_ID | Matrix_name |
|---|---|---|---|
| **1** | **0,0080** | **MA0144.1** | **STAT3** |
| 2 | 0,0134 | MA0058.1 | MAX |
| **3** | **0,0224** | **MA0137.2** | **STAT1** |
| 4 | 0,0278 | MA0093.1 | USF1 |
| 5 | 0,0317 | MA0258.1 | ESR2 |
| 6 | 0,0384 | MA0004.1 | Arnt |
| 7 | 0,0489 | MA0150.1 | NFE2L2 |

**Figure S11. ACE2 promoter Transcription factor binding motifs analysis.** TRAP analysis of transcription factors binding motifs in 1000 bp upstream sequence from ACE2 TSS (ACE2 proximal promoter). Analysis of TFs binding motifs using TRANSFAC database (**A**) and JASPAR database (**B**), alongside with related tables reporting ranking, p-values, TF matrix ID and TF binding motifs official name. STATs binding motifs are highlighted in different colours within ACE2 promoter sequences and reported in bold in the related table.

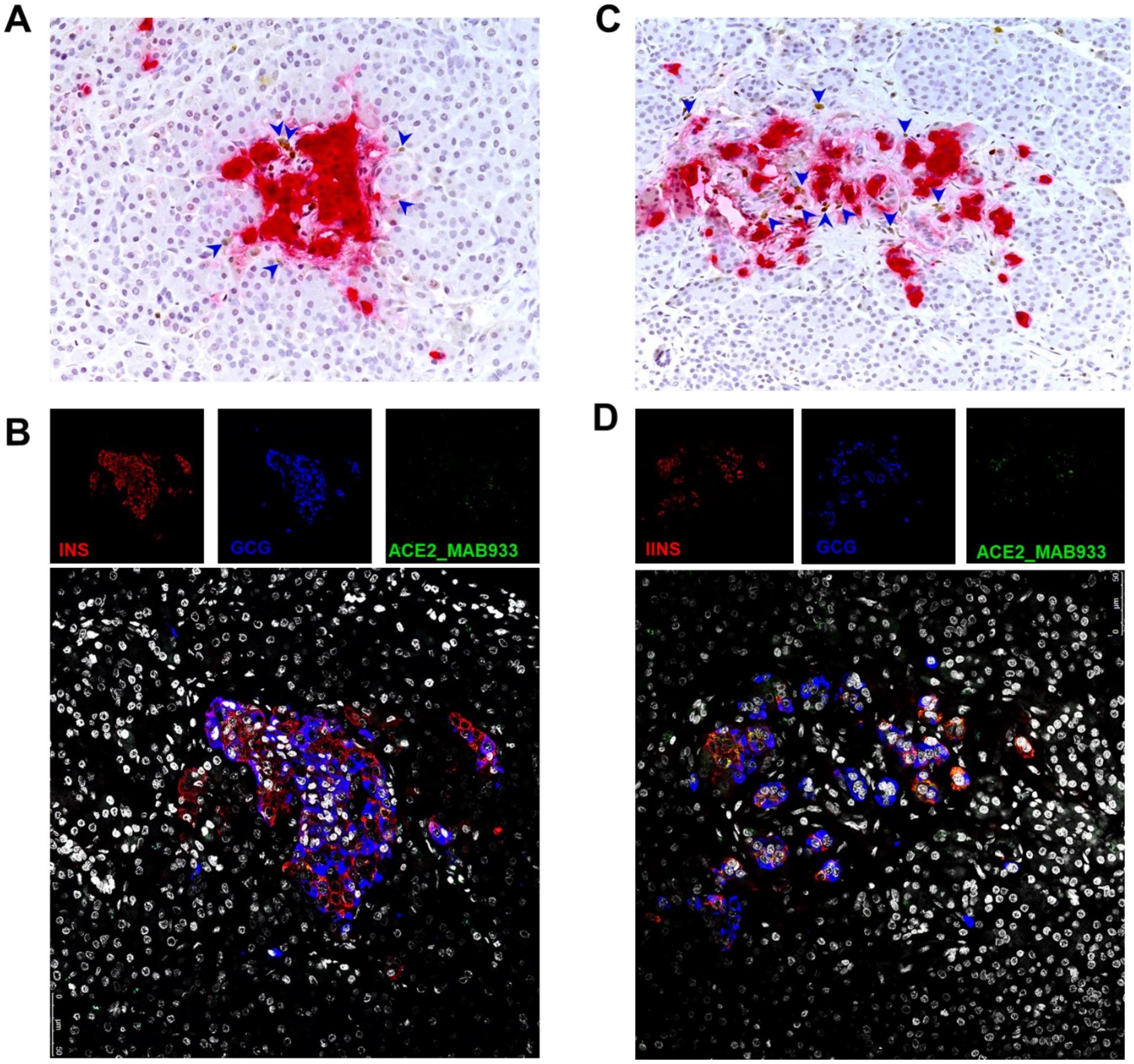

**Figure S12.** Triple immunofluorescence analysis of insulin (red), glucagon (blue) and ACE2 MAB933 (green) in a FFPE section of a longstanding T1D donor. Two infiltrated islets (A-B and C-D) were identified based on IHC insulin (red)-CD45 (brown) staining in a previous serial sections of the same T1D case.

**Table S1. Characteristics of non-diabetic multiorgan donors**. Main descriptive characteristics are reported alongside with tissue type obtained. Human islets (HI) purity values (only for pancreatic islets enzymatic isolation) and type of analysis performed, are reported as well.

| Case/ Sample ID | Gender | Age (years) | BMI (Kg/m2) | AutoAb (ELISA) | HiRes HLA | Ischemia Time (hours) | Cause of Death | Tissue type | HI Purity (Dithizone Stain- %) | HI Purity (insulin ICC-%) | Used for |
|---|---|---|---|---|---|---|---|---|---|---|---|
| 141117 CTR | M | 49 | 25.8 | GADA neg, IA-2A neg, ZnT8A neg | HLA:A*03,68; B*35,47; C*04,06; DRB1*03,08; DQB1*02,04 | 16h | Cardiovascular disease | FFPEand fresh-frozen pancreatic tissue | -- | -- | IHC/LCM |
| 181117 CTR | M | 39 | 23.6 | GADA neg, IA-2A neg ZnT8A neg | HLA:A*03,33; B*14, B*14 SD B65; C*08; DRB1*01, DQB1*05 | 18h | Trauma | FFPE and fresh-frozen pancreatic tissue | -- | -- | IHC/LCM |
| 110118 CTR | F | 46 | 32.5 | GADA neg IA-2A neg, ZnT8A neg | HLA:A*24; B*15,18; C*7; DRB1*04,11; DQB1*3 | 14h | Cardiovascular disease | FFPE and fresh-frozen pancreatic tissue | -- | -- | IHC/LCM |
| 300318 CTR | F | 56 | 24.5 | GADA neg, IA-2A neg, ZnT8A neg | HLA:A*2; B*07,50; C*06,07; DRB1*03,15; DQB1*02,06 | 17h | Trauma | FFPE and fresh-frozen pancreatic tissue | -- | -- | IHC |
| 210518 CTR | M | 44 | 24.5 | GADA neg, IA-2A neg, ZnT8A neg | HLA:A*24,34; B*15,55; C*03,07; DRB1*11,13; DQB1*3 | 36 | Cardiovascular disease | FFPE and fresh-frozen pancreatic tissue | -- | -- | IHC |
| 161118 CTR | F | 39 | 22.3 | GADA neg, IA-2A neg, ZnT8A neg | HLA:A LR*03,68; B*07,51; C*07,14; DRB1*01,15; DQB1*05,06 | 12h | Cardiovascular disease | FFPE and fresh-frozen pancreatic tissue | -- | -- | IHC/LCM |
| 110119 CTR | M | 22 | 27.2 | GADA neg, IA-2A neg, ZnT8A neg | HLA:A*01,02; B*08,18; C*07,07; DRB1*03,11; DQB1*02,03 | 42h | Trauma | FFPE and fresh-frozen pancreatic tissue | -- | -- | IHC/LCM |
| 060217 | F | 39 T1D duration :21 | 24,5 | GADA pos, IA-2A neg, ZnT8A neg | HLA:A*01,02; B*08,B*50; C*07, C*12; DRB1*03, DQRB1*17, DQB1*02 | 43h | Trauma | FFPE and fresh-frozen pancreatic tissue | -- | -- | IHC |
| Islet prep 1 | F | 23 | 22.5 | n/a | n/a | -- | Cardiac arrest | Enzymatic-isolated pancreatic islets | -- | 46 | RNA-seq (IL-1β + IFNγ treatment experiments) |
| Islet prep 2 | M | 31 | 27.8 | n/a | n/a | -- | Cerebral hemorrhage | Enzymatic-isolated pancreatic islets | -- | 66 | RNA-seq (IL-1β + IFNγ treatment experiments) |
| Islet prep 3 | M | 77 | 24.5 | n/a | n/a | -- | Cerebral hemorrhage | Enzymatic-isolated pancreatic islets | -- | 59 | RNA-seq (IL-1β + IFNγ treatment experiments) |
| Islet prep 4 | F | 64 | 29.4 | n/a | n/a | -- | Cerebral hemorrhage | Enzymatic-isolated pancreatic islets | -- | 47 | RNA-seq (IL-1β + IFNγ treatment experiments) |
| Islet prep 5 | F | 58 | 21.3 | n/a | n/a | -- | Cerebral hemorrhage | Enzymatic-isolated pancreatic islets | -- | 67 | RNA-seq (IL-1β + IFNγ treatment experiments) |

| | | | | | | | | | | | |
|---|---|---|---|---|---|---|---|---|---|---|---|
| Islet prep 6 | M | 67 | 25.7 | n/a | n/a | -- | Trauma | Enzymatic-isolated pancreatic islets | -- | 48 | RNA-seq (IFNα treatment experiments) |
| Islet prep 7 | F | 87 | 23.8 | n/a | n/a | -- | Cerebral hemorrhage | Enzymatic-isolated pancreatic islets | -- | 60 | RNA-seq (IFNα treatment experiments) |
| Islet prep 8 | F | 67 | 24.6 | n/a | n/a | -- | Cerebral hemorrhage | Enzymatic-isolated pancreatic islets | -- | 44 | RNA-seq (IFNα treatment experiments) |
| Islet prep 9 | F | 83 | 37.1 | n/a | n/a | -- | CVD | Enzymatic-isolated pancreatic islets | -- | 51 | RNA-seq (IFNα treatment experiments) |
| Islet prep 10 | F | 84 | 24.5 | n/a | n/a | -- | CVD | Enzymatic-isolated pancreatic islets | -- | 58 | RNA-seq (IFNα treatment experiments) |
| Islet prep 11 | F | 40 | 22.5 | n/a | n/a | -- | Stroke | Enzymatic-isolated pancreatic islets | -- | 59 | RNA-seq (IFNα treatment experiments) |
| Hi#2 | M | 52 | 34.2 | n/a | n/a | -- | n/a | Enzymatic-isolated pancreatic islets | ≥70% | | qRT-PCR ACE2 mRNA |
| Hi#3 | M | 50 | 27.4 | n/a | n/a | -- | n/a | Enzymatic-isolated pancreatic islets | ≥70% | | qRT-PCR ACE2 mRNA |
| Hi#4 | M | 55 | 28 | n/a | n/a | -- | n/a | Enzymatic-isolated pancreatic islets | ≥70% | | qRT-PCR ACE2 mRNA |
| Hi#8 | M | 38 | 23.1 | n/a | n/a | -- | Cardiovascular disease | Enzymatic-isolated pancreatic islets | 90% | | qRT-PCR ACE2 mRNA |

**Table S2.** ACE2-Insulin and ACE2-Glucagon colocalization rate (reported as percentage values) in human pancreatic islets of EUnPOD non-diabetic cases. Two different FFPE pancreas blocks per case were analysed; results for each block are reported separately in the table below.

| Case ID | Pancreas Block ID | ACE2-INS (Colocalization Rate- %) | ACE2-GCG (Colocalization Rate- %) |
|---|---|---|---|
| 110118 | Body 01A | 47.5% | 8.3% |
| 110118 | Head 02A | 59.3% | 12.2% |
| 110119 | Body 01A | 63.2% | 3.1% |
| 110119 | Tail 01A | 59.3% | 2.3% |
| 141117 | Body 01B | 83.6% | 6.6% |
| 141117 | Tail 03B | 55.0% | 5.6% |
| 161118 | Body 01A | 73.9% | 6.8% |
| 161118 | Tail 01A | 81.9% | 12.6% |
| 181117 | Head 02B | 53.6% | 7.0% |
| 181117 | Tail 01A | 62.2% | 9.2% |
| 210518 | Body 01B | 39.5% | 7.5% |
| 210518 | Head 02A | 35.6% | 11.3% |
| 300318 | Body 01A | 61.8% | 4.1% |
| 300318 | Tail 01A | 39.0% | 1.5% |

**Table S3.** Expression of ACE2 mRNA (ENST00000252519.8) in RNA-seq dataset of five different samples of EndoC-βH1, and five different Human Pancreatic islets preparations exposed or not to IL-1β + IFNγ (48h), and five different samples of EndoC-βH1 and six of Human Pancreatic islets exposed or not to IFNα (18h).

| **IL-1β + IFNγ (48h)** | | | | |
|---|---|---|---|---|
| | Not treated | Treated (IL-1β + IFNγ) | Log2 Fold Change (Cyt vs. nt) | pValue Adj |
| EndoC-βH1 | 0.061 | 1.05 | 4.19 | 3.02E-35 |
| Human Pancreatic islets | 2.20 | 5.76 | 1.51 | 7.902E-17 |
| **IFNα (18h)** | | | | |
| | Not treated | Treated (IFNα) | Log2 Fold Change (Cyt vs. nt) | pValue Adj |
| EndoC-βH1 | 0.057 | 3.15 | 5.92 | 9.35E-42 |
| Human Pancreatic islets | 3.17 | 16.30 | 2.39 | 8.45E-26 |

**Reagents Table**. Table reporting reagents or resources used in this study.

| REAGENT or RESOURCE | SOURCE | IDENTIFIER |
|---|---|---|
| ***Antibodies and reagent for IHC/IFA*** | | |
| Monoclonal Mouse anti-Human ACE2 | R&D System | MAB933, RRID:AB_2223153 |
| Monoclonal Rabbit anti-Human ACE2 | Abcam | Ab108252 RRID:AB_10864415 |
| Polyclonal Rabbit anti-Human ACE2 | Abcam | Ab15348 RRID:AB_301861 |
| Polyclonal Rabbit anti-Human CD31 | Abcam | Ab28364 RRID:AB_726362 |
| Polyclonal Rabbit Anti-Human Glucagon | Agilent/Dako | A0565 RRID:AB_10013726 |
| Monoclonal Mouse Anti-Human Glucagon | R&D System | MAB1249 RRID:AB_2107340 |
| FLEX Polyclonal Guinea Pig Anti-Insulin, Ready-to-Use | Agilent/Dako | IR002 RRID:AB_2800361 |
| Polyclonal Rabbit Anti-Mouse/HRP | Agilent/Dako | P0260 RRID:AB_2636929 |
| Polyclonal Goat Anti-Rabbit/HRP | Jackson ImmunoResearch Labs | 111-036-003 RRID:AB_2337942 |
| Goat Anti-Mouse IgG (H+L) Highly Cross-adsorbed Antibody, Alexa Fluor 488 | Thermo Fisher Scientific | A-11029 RRID:AB_2534088 |
| Goat anti-Rabbit IgG (H+L) Highly Cross-Adsorbed Antibody, Alexa Fluor 488 | Thermo Fisher Scientific | A-11034 RRID:AB_2576217 |
| Goat anti-Guinea Pig IgG (H+L) Highly Cross-Adsorbed Antibody, Alexa Fluor 555 | Thermo Fisher Scientific | A-21435 RRID:AB_2535856 |
| Goat anti-Guinea Pig IgG (H+L) Highly Cross-Adsorbed Antibody, Alexa Fluor 594 | Thermo Fisher Scientific | A-11076 RRID:AB_2534120 |
| Goat anti-Rabbit IgG (H+L) Highly Cross-Adsorbed Antibody, Alexa Fluor 594 | Thermo Fisher Scientific | A-11037 RRID:AB_2534095 |
| Goat anti-Rabbit IgG (H+L) Highly Cross-Adsorbed Antibody, Alexa Fluor 647 | Thermo Fisher Scientific | A-21245 RRID:AB_2535813 |
| Goat anti-Mouse IgG (H+L) Highly Cross-Adsorbed Antibody, Alexa Fluor 647 | Thermo Fisher Scientific | A-21236 RRID:AB_2535805 |
| ***Biological Samples*** | | |
| Human pancreatic islets | Tebu Bio/ University of Pisa | N/A |
| Human pancreatic tissues | EUnPOD INNODIA | N/A |
| Human lung tissues | University of Siena | N/A |
| ***Chemicals*** | | |
| Absolute Ethanol | VWR | 1.009.862.500 |
| Xilene | Merck | 28975325 |
| Hematoxylin | Sigma Aldrich | MHS31 |
| β-Mercaptoethanol | Sigma Aldrich | M7522 |
| LCM caps | Thermo Fisher Scientific | LCM0214 |
| DAPI | Sigma Aldrich | D8517 |
| Citrate buffer:<br>0.1 M Citric Acid Monohydrate<br>0.1M Sodium Citrate Tribasic Dihydrate | <br>Sigma Aldrich<br>Sigma Aldrich | <br>C1909<br>S4641 |
| ***Critical Commercial Assays*** | | |
| PicoPure RNA isolation kit Arcturus | Thermo Fisher Scientific | kit0204 |
| Agilent 2100 Bioanalyzer technology with RNA Pico chips | Agilent Technologies | 5067-1513 |
| SuperScript™ VILO™ cDNA Synthesis Kit | Thermo Fisher Scientific | 11754050 |

| | | |
|---|---|---|
| SensiFast Probe Lo-ROX Kit | Aurogene s.r.l | BIO-84020 |
| Direct-zol RNA Miniprep Kit | Zymo Research | R202 |
| Taq Man Preamp Master mix | Thermo Fisher Scientific | 4488593 |
| RNase-Free DNase Set | Qiagen | 79254 |
| ***Deposited Data*** | | |
| RNA-seq human islets and EndoC-βH1 exposed to IFNα | N/A | GSE133221 |
| RNA-seq human islets exposed to IL-1 β + IFNγ | N/A | GSE108413 |
| RNA-seq Endoc-βH1 exposed to IL-1 β + IFNγ | N/A | GSE137136 |
| ***Experimental Models: Cell Lines*** | | |
| EndoC-βH1 human β-cell line | UniverCell-Biosolutions (Toulouse-France) | N/A |
| ***Oligonucleotides*** | | |
| Endogenous control human GAPDH | Thermo Fisher Scientific | 4333764 |
| Endogenous control human ACTB | Thermo Fisher Scientific | 4333762 |
| Taq Man gene expression assay ACE2 | Thermo Fisher Scientific | Hs01085333 |
| ***Software and Algorithms*** | | |
| Expression Suite software 1.0.1 | Thermo Fisher Scientific | N/A |
| Leica TCS SP5 confocal laser scanning microscope system | Leica Microsystems | N/A |
| Laser Advanced fluorescence | Leica Microsystems | N/A |
| Opera Phenix High Content Screening System | PerkinElmer | N/A |
| Harmony® Office | PerkinElmer | N/A |
| Graph Pad Prism 8 | Prism | N/A |
| Qubit 3000 Fluorometer | Thermo Fisher Scientific | N/A |
| ***Other*** | | |
| DMEM high-glucose | Sigma Aldrich | 51441C |
| Penicillin/Streptomycin | Sigma Aldrich | P0781 |
| ECM gel from Engelbreth-Holm swarm murine sarcoma | Sigma Aldrich | E1270 |
| Fibronectin from bovine plasma | Sigma Aldrich | F1141 |
| DMEM low-glucose | Sigma Aldrich | D6046 |
| BSA fraction V | Sigma Aldrich | 10775835001 |
| L-Glutamine | Sigma Aldrich | G7513 |
| Penicillin/Streptomycin | Sigma Aldrich | P0781 |
| Nicotinamide | Sigma Aldrich | N0636 |
| Transferrin | Sigma Aldrich | T8158 |
| Sodium selenite | Sigma Aldrich | S5261 |
| Sodium Palmitate | Sigma Aldrich | P9767-5G |
| IL-1β | R&D System | 201-LB-005 |
| TNFα | Sigma Aldrich | T7539 |
| IFNγ | Roche | 11040596001 |
| CMRL culture medium | Thermo Fisher Scientific | 11-530-037 |
| Antibiotic/Antimicotic | Sigma Aldrich | A5955-100ML |
| Fetal Bovine Serum | Euroclone | EUS031153 |
| EUKITT mounting medium | Bio Optica | S9-25-37 |
| DAKO PAP pen | Agilent/Dako | S2002 |
| PBS w/o Ca2+ Mg2+ | Gibco | 14040-091 |
| Hydrogen peroxide | Sigma Aldrich | H1009 |

| | | |
|---|---|---|
| Bovine Serum Albumin | Sigma Aldrich | A1470-25G |
| Glycine | Thermo Fisher Scientific | 100006373 |
| DAB Novolink MAX DAB | Leica | RE7270-K |
| Vectashield | Vector Laboratories | H-1000 |
| Rabbit serum | Sigma Aldrich | R9133 |
| Goat serum | Sigma Aldrich | G9023 |
